# An *APOC1*^+^ inflammatory CAF-like state drives a senescent, treatment-resistant niche in rheumatoid arthritis

**DOI:** 10.64898/2026.04.17.718831

**Authors:** Risa Yoshihara, Sotaro Nakajima, Reo Yamazato, Tomoya Yoshida, Ikuo Takazawa, Yasunori Omata, Teh-Wei Wang, Kazuyoshi Ishigaki, Takahiro Itamiya, Mineto Ota, Yoichi Yasunaga, Yuichiro Fujieda, Takumi Matsumoto, Hirofumi Shoda, Kazuhiko Yamamoto, Naoto Tamura, Toshihide Mimura, Koichiro Ohmura, Akio Morinobu, Tatsuya Atsumi, Yoshiya Tanaka, Tsutomu Takeuchi, Yutaka Suzuki, Makoto Nakanishi, Tomohisa Okamura, Sakae Tanaka, Haruka Tsuchiya, Keishi Fujio

## Abstract

**Objectives:** Rheumatoid arthritis (RA) synovitis frequently persists despite cytokine-targeted therapies, suggesting the existence of pathogenic stromal programs that sustain chronic inflammation independently of canonical immune pathways. Although synovial fibroblasts (SF) are increasingly implicated in treatment resistance, the pathogenic fibroblast states driving refractory disease and their therapeutic vulnerabilities remain poorly defined.

**Methods:** We integrated multimodal single-cell and spatial profiling of synovial tissue from 54 patients with RA with prospective treatment-response data and functional studies in human fibroblasts and experimental arthritis models.

**Results:** We identified a *C-X-C motif chemokine 12 (CXCL12)*^hi^ *Apolipoprotein C1 (APOC1)*^+^ fibroblast population selectively enriched in treatment-refractory synovitis. Spatial analyses demonstrated that these fibroblasts establish CXCL12-dependent plasmablast niches within inflamed synovium, resembling inflammatory cancer-associated fibroblasts (iCAF) that orchestrate immune cell recruitment in the tumor microenvironment. *CXCL12*^hi^ *APOC1*^+^ fibroblasts exhibited a senescence-associated iCAF-like transcriptional program characterized by STAT3-C/EBP activation and APOC1 expression and were associated with poor response to TNF and IL-6 pathway inhibition. Mechanistically, *APOC1* knockdown in RA-SF attenuated invasive mesenchymal behavior and disrupted senescence-associated inflammatory programs, identifying APOC1 as a central regulator of pathogenic fibroblast reprogramming. Importantly, genetic or pharmacological elimination of senescent cells ameliorated experimental arthritis and enhanced the efficacy of TNF blockade.

**Conclusions:** These findings implicate iCAF-like fibroblasts with senescent properties as a mechanistic driver of refractory RA synovitis and highlight stromal senescence programs as preclinically actionable therapeutic targets beyond cytokine inhibition.

## Introduction

Rheumatoid arthritis (RA) is an autoimmune arthritis that causes perpetual joint damage and systemic inflammation, leading to physical dysfunction and shorter healthy life expectancy(*1*). Despite major advances in disease-modifying anti-rheumatic drugs (DMARDs) targeting inflammatory cytokine and signaling networks, 5–20% of patients remain refractory to successive therapies and are classified as difficult-to-treat (D2T) RA(*2*). Prolonged exposure to ineffective treatments imposes considerable individual and socioeconomic burdens, representing a major unmet clinical need(*2*).

Recent studies focusing on synovium, a primary target tissue in RA, have provided revolutionary insights into disease pathophysiology by adding information at local inflammatory sites. Synovial cellular composition varies markedly between patients and defines distinct inflammatory endotypes. In particular, mesenchymal cell-dominant phenotypes, especially enriched for synovial fibroblasts (SF), have been linked to resistance to cytokine-targeted therapies(*3–5*), suggesting that targeting non-immune stromal components could be promising for refractory cases. Although single-cell technologies have revealed cellular heterogeneity at unprecedented resolution, the responsible cell clusters and biological mechanisms defining RA endotypes remain incompletely elucidated.

Here, we established a multi-omics atlas of RA synovium incorporating single cell transcriptomics and proteomics, spatial profiles, and prospectively collected treatment response information. Integrative analyses identified a distinct cluster of SF that overexpressed *C-X-C motif chemokine 12* (*CXCL12*) and *Apolipoprotein C1* (*APOC1*) and was associated with resistance to cytokine blocking agents. This SF cluster exhibited tumor-like mesenchymal features alongside a feature of inflammatory cancer-associated fibroblast (iCAF) with senescence-associated secretory phenotype (SASP). Our findings suggest that therapy-resistant RA represents a fibroblast-driven pathological state with cancer-like stromal reprogramming.

## Results

### Refractory synovitis is dominated by synovial fibroblasts

We collected synovial samples from 54 active RA patients and constructed the following datasets: quantitative composition of major cell types (flow cytometry; FCM), single-cell transcriptome profiles, single-nucleus chromatin accessibility and transcriptome profiles, spatial transcriptomic (10x Genomics) and proteomic (PhenoCycler-Fusion system, Quanterix) maps, and prospective clinical information (figure 1A, supplemental figure S1 and supplemental table 1). We subsequently performed functional analyses to explore the molecular mechanisms underlying pathogenic cell states (figure 1, B to E).

**figure 1.**
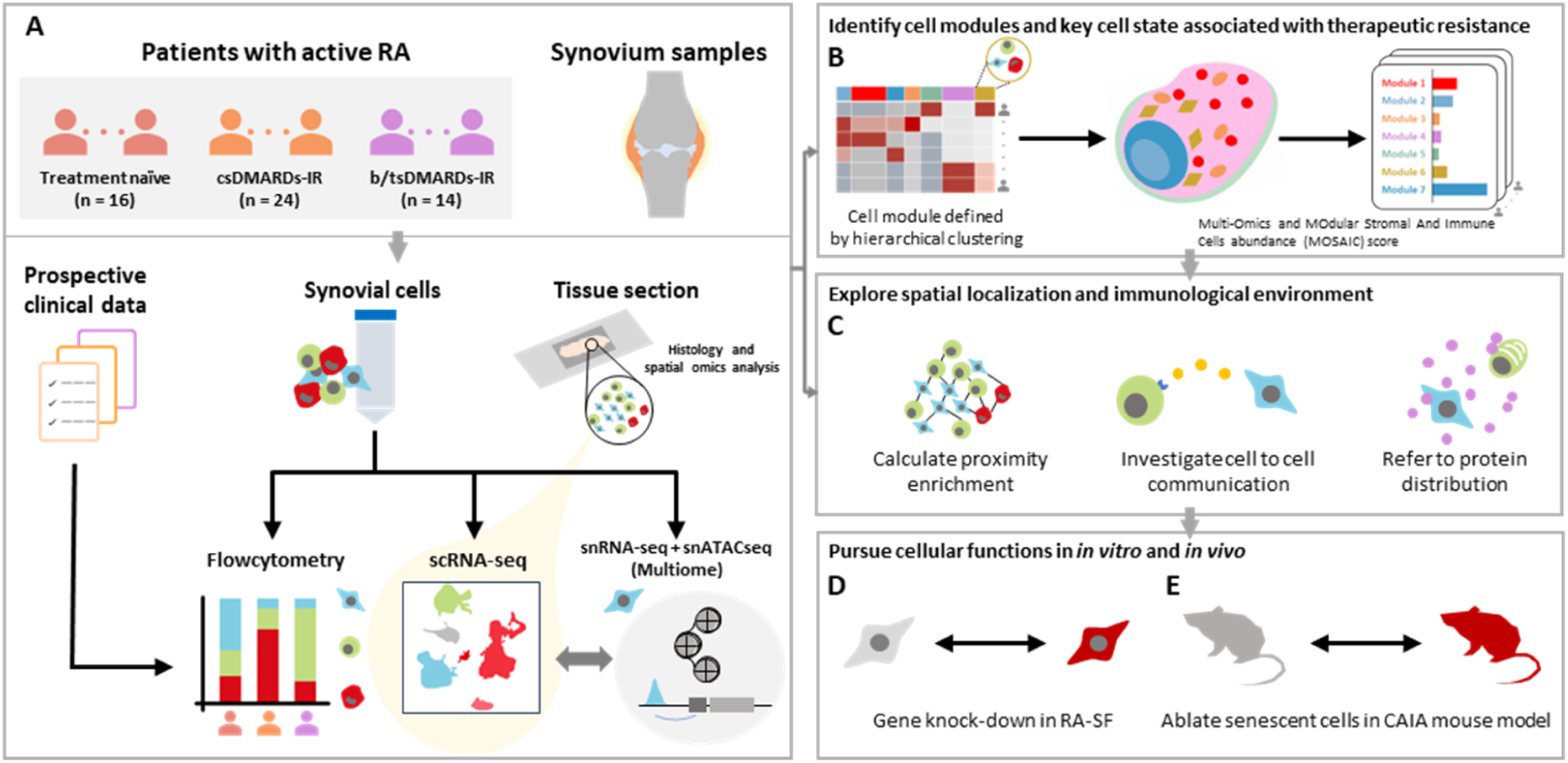
Overview of the multimodal approach that highlights the contribution of *CXCL12*^hi^ *APOC1*⁺ fibroblasts to refractory synovitis in rheumatoid arthritis. **(A)** Schematic overview of patient recruitment, prospective clinical assessment, and synovial tissue processing, including single-cell and single-nucleus profiling of the transcriptome and chromatin accessibility. **(B)** Stratification of synovial cell modules and calculation of the MOdular Stromal And Immune Cells (MOSAIC) score. **(C)** Evaluation of spatial organization and dynamics of the synovial microenvironment associated with treatment resistance. **(D and E)** Functional *in vitro* and *in vivo* assays using synovial fibroblasts (SF) from patients with rheumatoid arthritis (RA) **(D)** and experimental mouse models **(E)**. b/tsDMARDs, biologic and targeted synthetic disease-modifying antirheumatic drugs; CAIA, collagen antibody-induced arthritis; csDMARDs, conventional synthetic disease-modifying antirheumatic drugs; FCM, flow cytometry; IR, inadequate responder; MOSAIC, MOdular Stromal And Immune Cells abundance; RA, rheumatoid arthritis; RA-SF, rheumatoid arthritis synovial fibroblast; scRNA-seq, single-cell RNA sequencing; SF, synovial fibroblast; snATAC-seq, single-nucleus assay for transposase-accessible chromatin sequencing; snRNA-seq, single-nucleus RNA sequencing.

To identify cell populations that persist in inflamed synovium despite treatment, we examined how synovial composition differs by treatment status. We evaluated the proportion of CD4^+^ T cells, CD8^+^ T cells, CD19^+^ B cells, monocytes, natural killer (NK) cells, SF, vascular endothelial cells (ECs) and mural cells by FCM analysis (supplemental figure S2). Of note, with more intensive treatment—methotrexate (MTX) or biological or targeted synthetic DMARDs (b/tsDMARDs)—mesenchymal cells, especially SF, became more dominant than immune cells (supplemental figure S3A). We applied multivariate linear regression to examine how each medication affects cell proportions, adjusting for CDAI (Clinical Disease Activity Index) and the method of sample collection. As a result, MTX (supplemental figure S3B) and prednisolone (PSL) (supplemental figure S3C) were associated with reduced CD8^+^ and CD4^+^ T-cell infiltration, respectively, while b/tsDMARDs had limited effects on immune cell composition (supplemental figure S3D). In contrast, the proportion of SF was significantly increased with the use of MTX and PSL. Recently, Cell Type Abundance Phenotypes (CTAPs) were proposed based on the frequency of major cell types(*4*). CTAP-F (SF-dominant) accounted for 53% of patients after treatment with bDMARDs, and CTAP-EFM (EC-, SF-, and myeloid-dominant) included more TNF-inadequate responders than immune cell-dominant types(*4*). Thus, antirheumatic drugs have a significant impact on synovial cells composition, and remaining SF may maintain and amplify inflammation in refractory cases.

### Synovial single-cell atlas provides 26 major cell populations and 10 fine SF clusters

To gain insights into intractable disease, we constructed a synovial single-cell atlas of 82,708 cells from 54 patients. We identified 26 major cell populations (17 hematopoietic, 9 stromal), using Uniform Manifold Approximation and Projection (UMAP) (supplemental figure S4, A to D).

Hematopoietic cells comprised 4 CD4^+^ T cell, 3 CD8^+^ T cell, 4 macrophage, 2 NK cell, 2 dendritic cell (DC) clusters, along with B cells and plasmablasts. CD4^+^ T cells included naïve and two effector-memory subsets, *FOXP3*^+^ regulatory T cells, follicular helper T cells (Tfh) and peripheral helper T (Tph)-like cells. CD8^+^ T cells were classified by cytotoxic molecule expression into: *GZMK*⁺ *GZMB*⁺ CD8⁺ T cells, which expand clonally in inflamed joints and can drive tissue inflammation via IFN-γ and activating complement(*6*), mucosal-associated invariant T (MAIT) cells, and cytotoxic T lymphocytes (CTL). Macrophages comprised *SPP1*^+^ macrophages, *MERTK*^+^ macrophages—which exert homeostatic and regulatory functions via the pro-resolving mediator resolvin D1 and promote repair responses in SF(*7*)—along with *CLEC10A*^+^ macrophages and *S100A12*^+^ macrophages. NK cells were split by *CD56* and *CD16* expression into *CD56*^dim^ *CD16*^+^ and *CD56*^br^ *CD16*^−^ NK cells(*4*). Two DC clusters, plasmacytoid and conventional, were distinctly separated(*8*).

Stromal cells comprised five SF, three EC, and one mural cell cluster. SF included four sublining SF (*THY1*^hi^, *THY1*^low^, *CD34*^+^ *THY1*^hi^ and *CD34*^+^ *THY1*^low^) and one lining SF(*9, 10*). Five major SF clusters were subdivided into 10 fine clusters (figure 2A and supplemental figure S4, E and F). Three subclusters of *HLA-DR*^hi^ sublining SF—a major producer of *IL6* and an expanded population in RA synovium relative to osteoarthritis—were identified: *IL6*^+^ *HLA-DR*^hi^ sublining expressing *NOTCH3*, *CHI3L2*^hi^ *HLA-DR*^hi^ sublining, and *CD74*^hi^ *HLA-DR*^hi^ sublining. *POSTN*^+^ sublining, which distributes around blood vessels(*11*), expressed angiogenic factors such as *MDK* and *PTN*. *CXCL12*^hi^ *APOC1*^+^ sublining exhibited relatively high *CXCL12* expression, a potent chemotactic factor that recruits immune cells and contributes to angiogenesis and bone destruction(*12*). It also characteristically expressed lipid-processing molecules including *APOC1* and *APOE*(*13*). *APOD*^+^ sublining expressed *GAS6*, possibly contributing to the homeostatic regulatory function of *MERTK*^+^ macrophages(*7*). *DKK3*^+^ sublining and *CD34*^+^ sublining (*THY1*^hi^ and *THY1*^low^) were also identified; the latter highly expressed *PI16* and *DPP4*, representative molecules of pan-tissue fibroblasts(*14*). *PRG4*^+^ lining distinctively expressed an extracellular protease, *MMP3*. Aside from SF, *APLN*^+^ capillary EC, *ACKR1*^+^ venular EC, arterial EC, and mural cells were distinguished.

**figure 2.**
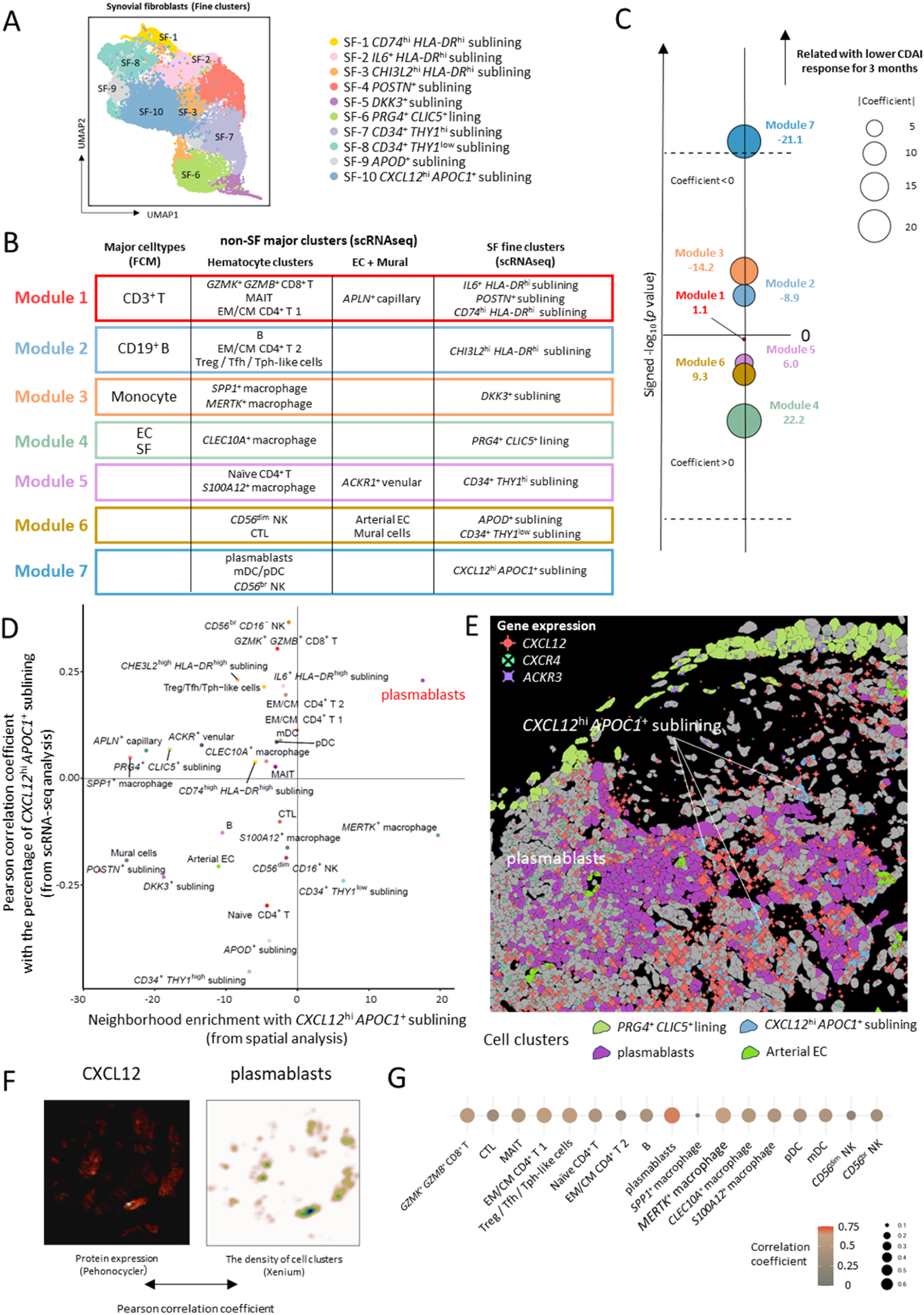
*CXCL12*^hi^ *APOC1*^+^ fibroblasts and plasmablasts form a microenvironment that promotes refractory synovitis. **(A)** Identification of 10 synovial fibroblast (SF) subclusters, colored by fine-grained cellular states. **(B)** Summary of synovial cell modules defined by hierarchical clustering (supplemental figure S5). **(C)** Association of the MOdular Stromal And Immune Cells (MOSAIC) score with reduction in the Clinical Disease Activity Index (CDAI) over 3 months following treatment with cytokine-blocking agents (tumor necrosis factor (TNF) inhibitors or interleukin-6 receptor (IL-6R) inhibitors) administered after synovial biopsy. Bubble size, the absolute value of coefficients derived from multivariate linear regression analysis; black dashed lines, Bonferroni-adjusted significance thresholds. **(D)** Quantitative correlation in scRNA-seq analysis and neighborhood enrichment calculated by spatial analysis between *CXCL12*^hi^ *APOC1*^+^ fibroblasts and other synovial cell clusters. **(E)** Spatial distribution of *CXCL12*^hi^ *APOC1*^+^ fibroblasts and plasmablasts in synovial tissue from a patient with active rheumatoid arthritis, together with the expression of *CXCL12*, *CXCR4* and *ACKR3*. **(F)** Comparison of kernel density estimates of hematopoietic cell clusters, represented by plasmablasts, and CXCL12 protein distribution. **(G)** Quantitative correlation between plasmablast density and CXCL12 protein abundance. Bubble size, the absolute value of coefficients calculated using Pearson’s correlation analysis. CDAI, Clinical Disease Activity Index; CTL, cytotoxic T lymphocyte; EC, endothelial cell; IL-6R, interleukin-6 receptor; MAIT, mucosal-associated invariant T cell; mDC, myeloid dendritic cell; MOSAIC, MOdular Stromal And Immune Cells abundance; NK, natural killer cell; pDC, plasmacytoid dendritic cell; RA, rheumatoid arthritis; SF, synovial fibroblast; Tfh, follicular helper T cell; TNF, tumor necrosis factor; Tph, peripheral helper T cell; Treg, regulatory T cell.

### Synovial cell modules are stratified by selectively enriched cell states

Recently, inter- and intra-individual diversity of local inflammation has drawn attention via deconstruction of RA synovium (*4, 10, 15*). To relate treatment response to cell clusters coexistence, we performed hierarchical clustering using: the percentage of major synovial cell types (CD3^+^ T cells, CD19^+^ B cells, monocytes, SF, and EC) among total live cells by FCM, of major hematopoietic cell clusters among total hematopoietic cells, major non-SF stromal cell clusters among total non-SF stromal cells, and SF fine clusters among total SF by scRNA-seq (figure 1B, see supplementary materials and methods). Consequently, we identified seven clusters of coexisting cell states, termed ‘synovial cell modules’ (figure 2B and supplemental figure S5). Briefly, module 1 was characterized by *GZMK*⁺ *GZMB*⁺ CD8⁺ T cells, and two *HLA-DR*⁺ sublining SF clusters (*IL6*^+^ *HLA-DR*^hi^ sublining and *CD74*^hi^ *HLA-DR*^hi^ sublining). Module 2 contained B cells and Tph/Tfh cells that help B cell activation and antibody production(*16*), suggesting the presence of tertiary lymphoid follicles. Module 3 was characterized by the coexistence of abundant myeloid cells (*MERTK*^+^ macrophages and *SPP1*^+^ macrophages) and *DKK3*^+^ sublining SF. Of note, a subpopulation of *DKK3*^+^ sublining SF coexpressing *CD200* has been reported to induce resolution of arthritis via CD200-CD200R1 signaling(*17*) and crosstalk with *MerTK*^+^ macrophages to resolve fibrosis(*18*). Module 4 was composed of *CD34*^+^ *THY1*^hi^ sublining SF, naïve CD4^+^ T cells and *S100A12*^+^ macrophages. Module 5 mainly consisted of stromal cells, particularly *PRG4*^+^ *CLIC5*^+^ lining SF, consistent with lining SF enrichment in CTAP-F, a SF-abundant phenotype(*4*). Module 6 demonstrated marked enrichment of mural cells and arterial EC. Module 7 featured *CXCL12*^hi^ *APOC1*^+^ sublining SF, plasmablasts, myeloid and plasmacytoid DCs.

Next, each module’s involvement per each patient was quantified by multidimensional scaling (MDS) and defined as the Multi-Omics and MOdular Stromal And Immune Cells abundance (MOSAIC) score (supplemental figure S6). Unlike classical approaches that assign each patient a single pathotype, the MOSAIC score simultaneously quantifies multiple pathological states within a patient. We related the seven modules’ MOSAIC scores (MOSAIC-m1 to MOSAIC-m7) to treatment response—CDAI reduction over 3 months in 29 patients who received IL-6R or TNF inhibitors after biopsy. Importantly, the MOSAIC-m7 score was significantly associated with lower CDAI response (figure 2C), indicating that the local tissue environment composed of *CXCL12*^hi^ *APOC1*^+^ sublining SF and coexisting immune cells (plasmablasts and DCs) contributes to intractable pathophysiology of RA.

### *CXCL12*^hi^ *APOC1*^+^ sublining SF forms a treatment-resistance niche with plasmablast infiltration

To further investigate local niches associated with MOSAIC-m7, we performed spatial multi-omics analysis on 5,101 transcripts and 60 proteins of synovial samples from 7 patients with RA (figure 1C). We annotated cell types through robust cell type deconvolution (RCTD) learned from our scRNA-seq dataset (supplemental figure S7, A and B, and supplemental figure S8). Next, we proceeded to explore the spatial proximity between *CXCL12*^hi^ *APOC1*^+^ sublining SF and hematopoietic cells or stromal cells by Giotto. Notably, *CXCL12*^hi^ *APOC1*^+^ sublining SF were spatially adjacent to each other and to plasmablasts, a common MOSAIC-m7 component, and the percentages of *CXCL12*^hi^ *APOC1*^+^ sublining SF and plasmablasts were also positively correlated in scRNA-seq (figure 2D). While *CXCL12*^hi^ *APOC1*^+^ sublining SF expresses high *CXCL12*, the inference of cell-to-cell communication by CellChat v2 revealed that *CXCL12*^hi^ *APOC1*^+^ sublining SF and plasmablasts communicate through the *CXCL12*-*CXCR4* axis. *CXCL12*^hi^ *APOC1*^+^ sublining SF also retrieves *CXCL12* by expressing *ACKR3,* a receptor of *CXCL12* (figure 2E and supplemental figure S9), indicating an endogenous activation pathway(*19*). Furthermore, because CXCL12 diffuses within tissues and supports B cell differentiation and plasmablast survival(*20*), we examined CXCL12 protein levels and the density of each immune cell type on the same tissue section (figure 2F). The density of plasmablast showed the strongest correlation with CXCL12 protein distribution among all hematopoietic clusters at single-cell resolution (figure 2G), consistent with a previous report showing high *CXCL12* RNA expression in the plasma cell-rich areas of inflamed synovial tissue(*21*). Our data extend the earlier observation that antigen-secreting cells are abundant in synovium from inadequate responders to TNF inhibitors(*22*).

### The property of *CXCL12*^hi^ *APOC1^+^* sublining SF cannot be easily modified qualitatively by current therapeutic agents

Our findings suggested that *CXCL12*^hi^ *APOC1*^+^ sublining SF forms a CXCL12-rich stromal niche which promotes plasmablasts accumulation even under cytokine inhibition. We next sought to characterize the influence of major proinflammatory cytokines on each SF fine cluster. A genome-wide transcript catalog of RA-SF stimulated with eight inflammatory cytokines (IFN-α, IFN-γ, TNF-α, IL-1β, IL-6/sIL-6R, TGF-β1, IL-17, IL-18) was previously constructed by us(*23*). We defined cytokine stimulation signatures as the top 1,000 genes (supplemental table 2) significantly upregulated by each individual cytokine, and applied gene set variation analysis (GSVA) to estimate the cytokine activity in each SF fine cluster (figure 3A). As a result, three clusters of *HLA-DR*^hi^ sublining SF (*IL6*^+^ *HLA-DR*^hi^ sublining, *CHI3L2*^hi^ *HLA-DR*^hi^ sublining and *CD74*^hi^ *HLA-DR*^hi^ sublining) were enriched for IFN-γ, TNF-α and IL-6 signatures (figure 3B), consistent with the reports that *HLA-DR*^hi^ sublining SF can be induced by IFN-γ and TNF-α from lymphocytes(*6, 10*). *PRG4*^+^ lining SF had enhanced IL-1β and TNF-α signatures, which may reflect its colocalization with IL-1β and TNF-α-producing tissue macrophages(*24*). *POSTN*^+^ sublining SF and *DKK3*^+^ sublining SF showed enhanced TGF-β signatures, both highly expressing myofibroblastic gene markers such as *ACTA2, SMAD2* and *PDGFRB* (supplemental figure S4F), in line with the previous report that TGF-β promotes myofibroblast differentiation and activation(*25*). Interestingly, *CXCL12*^hi^ *APOC1*^+^ sublining SF was distinctive in that it showed little enrichment of cytokine signatures, despite their presence in active synovitis.

**figure 3.**
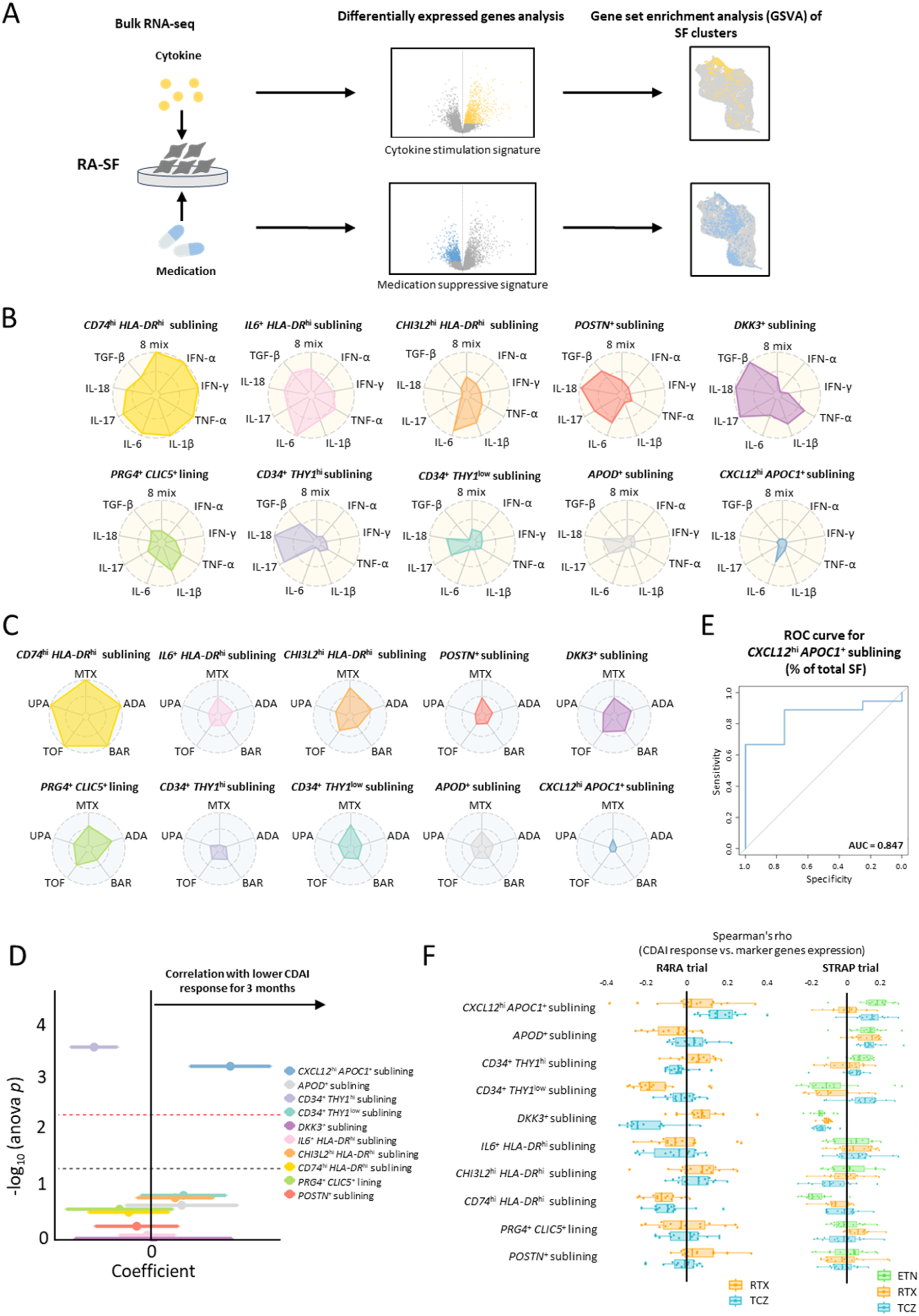
*CXCL1*2^hi^ *APOC1*^+^ fibroblasts are characterized by low proinflammatory cytokine activity and poor responses to cytokine-blocking therapies. **(A**) Schematic overview of the computational pipeline used to derive cytokine stimulation and medication suppressive signatures. Using a gene perturbation dataset of synovial fibroblasts (SF) exposed to eight inflammatory stimuli (TNF-α, IL-1β, IL-6, IFN-γ, IFN-α, IL-17, IL-18, and TGF-β) or five disease-modifying agents (methotrexate, adalimumab, baricitinib, tofacitinib, and upadacitinib), the top 1,000 regulated genes were defined as each signature. Signature scores were calculated using Gene Set Variation Analysis (GSVA). **(B and C)** GSVA scores of cytokine stimulation signatures (**B**) and medication suppressive signatures (**C**) across SF fine clusters. **(D)** Association between the abundance of each SF fine cluster and reduction in the Clinical Disease Activity Index (CDAI) over 3 months following treatment with cytokine-blocking agents (TNF inhibitors or interleukin-6 receptor (IL-6R) inhibitors) administered after synovial biopsy in 25 patients. Coefficients and association *p* values were calculated using mixed-effects association of single cells (MASC). Red and black dashed lines indicate Bonferroni-adjusted significance thresholds and *p* = 0.05, respectively. **(E)** A receiver operating characteristic (ROC) curve showing the performance of the proportion of *CXCL1*2^hi^ *APOC1*^+^ sublining among total SF for predicting failure to achieve a 50% improvement in CDAI with cytokine-blocking agents (TNF inhibitors or interleukin-6 receptor (IL-6R) inhibitors) in 22 patients (responders, n = 18; non-responders, n = 4). **(F)** Correlation between CDAI change at 16 weeks and expression of the top 15 differentially expressed genes (DEGs) defining SF fine clusters in publicly available bulk RNA-seq datasets from the R4RA trial (left) and the STRAP trial (right). Dots, Spearman’s rank correlation coefficients. ADA, adalimumab; BAR, baricitinib; CDAI, Clinical Disease Activity Index; DEGs, differentially expressed genes; ETN, etanercept; GSVA, Gene Set Variation Analysis; IFN, interferon; IL, interleukin; MASC,mixed-effects association of single cells; MTX, methotrexate; ROC, receiver operating characteristic ; R4RA, Rituximab versus tocilizumab in rheumatoid arthritis; SF, synovial fibroblast; STRAP, Stratification of Biologic Therapies for RA by Pathobiology; TCZ, tocilizumab; TGF, transforming growth factor; TNF, tumor necrosis factor; TOF, tofacitinib; UPA, upadacitinib; RTX, rituximab.

To assess the impact of existing disease-modifying agents, we employed a bulk RNA-seq database of RA-SF treated with MTX, a TNF inhibitor (adalimumab; ADA), and three JAK inhibitors (baricitinib; BARI, tofacitinib; TOFA, upadacitinib; UPA)(*26*). Each medication-suppressive signature score was calculated using the top 1,000 genes significantly down-regulated after 24-hour treatment by each agent (supplemental table 3). Consequently, *CD74*^hi^ *HLA-DR*^hi^ sublining SF with especially active IFN-γ, IL-6 and TNF signatures appeared to be transcriptionally regulated by MTX, JAK inhibitors, and a TNF inhibitor (figure 3C). Strikingly, we found that *CXCL12*^hi^ *APOC1*^+^ sublining SF scarcely displayed any medication-suppressive signatures. These results highlighted *CXCL12*^hi^ *APOC1*^+^ sublining SF as a notable cluster neither induced by major proinflammatory cytokines nor qualitatively modified by existing agents.

These results were validated when mixed-effects association testing for single cells (MASC) was applied to the abundance of each SF fine cluster and treatment response to cytokine-blocking agents in our cohort (figure 3D), and the proportion of *CXCL12*^hi^ *APOC1*^+^ sublining SF within the total SF showed a predictive ability for responses to IL-6R inhibitors or TNF inhibitors (CDAI ≥ 50% improvement), with an AUC of 0.847 (95% CI, 0.668–1.000) (figure 3E). Strikingly, in a larger public bulk RNA-seq dataset of synovial tissue from the R4RA (rituximab versus tocilizumab in anti-TNF inadequate responder patients with RA) trial (figure 3F), the first biopsy-based randomized clinical trial comparing tocilizumab (TCZ) with rituximab (RTX) in anti-TNF inadequate responders stratified by synovial B cell signatures(*27*), the expression levels of all top differentially expressed genes (DEGs) in *CXCL12*^hi^ *APOC1*^+^ sublining SF were positively correlated with CDAI changes after TCZ treatment (supplemental table 4), suggesting treatment resistance; this was statistically significant by a binomial test (*p* = 3.05×10^−5^). Similar results were consistently observed in TCZ (*p* = 7.39×10^−3^) and etanercept (ETN) (*p* = 9.77×10^−4^) cohorts from the STRAP (stratification of biologic therapies for RA by pathobiology) trial in biologic-naive patients(*28*) (figure 3F and supplemental table 5). These findings implicate *CXCL12*^hi^ *APOC1*^+^ sublining SF as a key driver of refractoriness to cytokine inhibitors.

### *CXCL12*^hi^ *APOC1*^+^ sublining SF parallels iCAF and exhibits tumor-like properties

Stromal remodeling in the RA synovium closely resembles that observed in cancer. RA-SF undergo tumor-like expansion within the inflamed synovium(*29*), mirroring CAF that accumulate in the tumor microenvironment (TME) and promote tumor progression through matrix production and inflammatory signaling(*30*). This analogy prompted us to investigate whether *CXCL12*^hi^ *APOC1*^+^ sublining SF shares functional properties with CAF. GSVA showed that *CXCL12*^hi^ *APOC1*^+^ sublining SF exhibits transcriptional features characteristic of iCAF described in breast and pancreatic cancers(*31–35*) (figure 4A and supplemental figure S10A). iCAF secrete inflammatory mediators and modulate immune cell recruitment in the TME, contributing to chemoresistance(*36*). In contrast, *IL6*^+^ *HLA-DR*^hi^ sublining SF showed greater transcriptional similarity to myofibroblastic CAF (myCAF), contributing to matrix deposition and tissue remodeling(*36*). *CD74*^hi^ *HLA-DR*^hi^ sublining SF resembled antigen-presenting CAF (apCAF), characterized by major histocompatibility complex (MHC) class II expression and antigen presentation to immune cells(*36*).

**figure 4.**
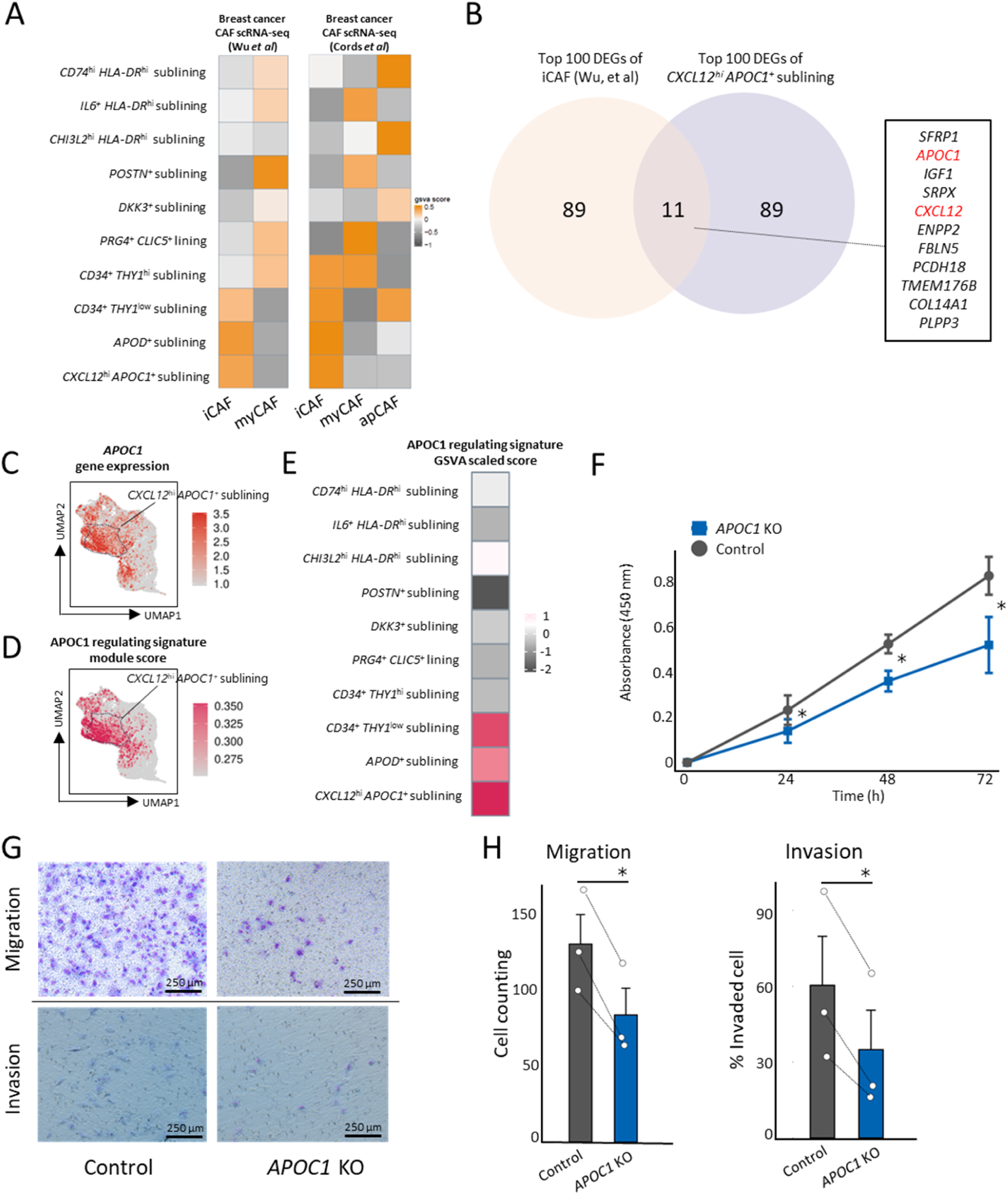
*APOC1* acts as a key regulator of tumor-like traits and *CXCL12* expression in synovial fibroblasts. **(A)** Heatmap showing Gene Set Variation Analysis (GSVA) scores of selected gene signatures derived from published cancer-associated fibroblast (CAF) populations in breast cancer across synovial fibroblast (SF) fine clusters. **(B)** Overlap between the top 100 differentially expressed genes (DEGs) of inflammatory CAF (iCAF) in breast cancer and *CXCL12*^hi^ *APOC1*^+^ sublining SF. The boxed panel shows all overlapping genes ranked by log fold change of the average expression between *CXCL12*^hi^ *APOC1*^+^ sublining SF and other SF clusters. **(C and D)** Concordance between *APOC1* expression (**C**) and the APOC1-regulated gene signature (**D**). Module scores represent the relative expression of gene sets altered in primary human rheumatoid arthritis synovial fibroblasts (RA-SF) following *APOC1* knockdown. **(E)** GSVA scores of the APOC1-regulated signature across SF fine clusters. **(F)** Cell viability assessed by Cell Counting Kit-8 (CCK-8) assays in MH7A cells depleted of *APOC1* (blue) or negative control (black). Horizontal crossbars, mean; error bars, SD. *P* values, a paired t-test from four independent experiments (* *p* <0.05). **(G and H)** Representative images (**G**) and quantification (**H**) of the migration and invasion activities of MH7A cells depleted of *APOC1* (blue) or negative control (black). Bars, mean; error bars, SD. *P* values, a paired t-test from three independent experiments (* *p* <0.05). apCAF, antigen-presenting cancer-associated fibroblast; CAF, cancer-associated fibroblast; CCK-8, Cell Counting Kit-8; DEGs, differentially expressed genes; GSVA, Gene Set Variation Analysis; iCAF, inflammatory cancer-associated fibroblast; KO, knockout; myCAF, myofibroblastic cancer-associated fibroblast; nm, nanometre; RA-SF, rheumatoid arthritis synovial fibroblasts; ROS, reactive oxygen species; scRNA-seq, single-cell RNA sequencing; SD, standard deviation; SEM, standard error of the mean; SF, synovial fibroblast.

Several scRNA-seq studies have reported that lipid-processing genes are enriched in iCAF signatures (*31–33*). Consistently, *APOC1*—an apolipoprotein C family member central to lipid transport and metabolism—(*13*)emerged as one of the top DEGs in *CXCL12*^hi^ *APOC1*^+^ sublining SF and overlaps with iCAF signatures identified in breast cancer(*31*) (figure 4B and supplemental table 6 and 7). APOC1 functions as an oncogenic factor that promotes cancer cell proliferation, invasion and metastasis through epithelial-mesenchymal transition (EMT), MAPK/ERK and STAT3 signalling(*37*). Elevated APOC1 expression is associated with poor prognosis in multiple malignancies(*38*). We generated *APOC1* knock-down cells from a human primary RA-SF using CRISPR–Cas9 (figure 1D) and performed bulk RNA-seq (supplemental table 8). Gene set enrichment analysis (GSEA) revealed that genes upregulated in iCAF(*31, 32*) (supplemental figure S10B) were significantly enriched among genes downregulated following *APOC1* knock-down in RA-SF. Furthermore, *APOC1* knock-down resulted in coordinated alterations in the expression of *APOC1* itself (figure 4C) and genes highly expressed by *CXCL12*^hi^ *APOC1*^+^ sublining SF (figure 4D). Consistently, *CXCL12*^hi^ *APOC1*^+^ sublining SF exhibited the strongest *APOC1*-regulated transcriptional signature among SF fine clusters (figure 4E), indicating that *APOC1* is an upstream determinant of this phenotype. *APOC1* knock-down significantly reduced the viability (figure 4F), migration and invasion (figure 4G,H) of MH7A cells, a human RA-SF cell line. Collectively, these results identify *APOC1* as a central regulator that confers *CXCL12*^hi^ *APOC1*^+^ sublining SF with both iCAF-like transcriptional features and tumor-like properties.

### STAT3-C/EBP cascade governs the transcriptional program of *CXCL12*^hi^ *APOC1*^+^ sublining SF

To define the transcriptional circuitry underlying *CXCL12*^hi^ *APOC1*^+^ sublining SF, we integrated single-nucleus transcriptomic and chromatin accessibility profiling from SF of nine patients with active RA with our scRNA-seq dataset. Chromatin-based clustering and transcriptomic projection identified ATAC_SF2 as the chromatin-defined counterpart of *CXCL12*^hi^ *APOC1*^+^ sublining SF (figure 5A and supplemental figure S11, A and B). Notably, ATAC_SF2 exhibited marked enrichment of motifs for C/EBP family and STAT3 (figure 5B), implicating them as candidate upstream regulators. Peak-to-gene analysis indicated that binding of C/EBP members and STAT3 to regulatory regions surrounding *CXCL12* is associated with its transcriptional activation in ATAC_SF2 (figure 5C and supplemental figure S11C). Moreover, consistent with observations in mammary epithelial cells and hepatocytes, STAT3 occupancy near the *CEBPB* locus correlated with *CEBPB* expression (supplemental figure S11D). Given that STAT3 has been reported to directly interact with APOC1 in multiple cellular contexts(*37*), *APOC1* knockdown reduced STAT3 phosphorylation in RA-SF (figure 5D), supporting the involvement of an APOC1–STAT3–C/EBP axis in regulating CXCL12 expression in *CXCL12*^hi^ *APOC1*^+^ sublining SF. Consistently, analysis of a fibroblast CRISPR activation (CRISPRa) Perturb-seq dataset(*39*) demonstrated that CEBPB activation induces the core transcriptional program of *CXCL12*^hi^ *APOC1*^+^ sublining SF, including *CXCL12* expression (figure 5E). These observations align with reports identifying the CEBP family as key drivers of iCAF phenotypes(*40*), suggesting shared transcriptional regulatory mechanisms between iCAF and *CXCL12*^hi^ *APOC1*^+^ sublining SF.

**figure 5.**
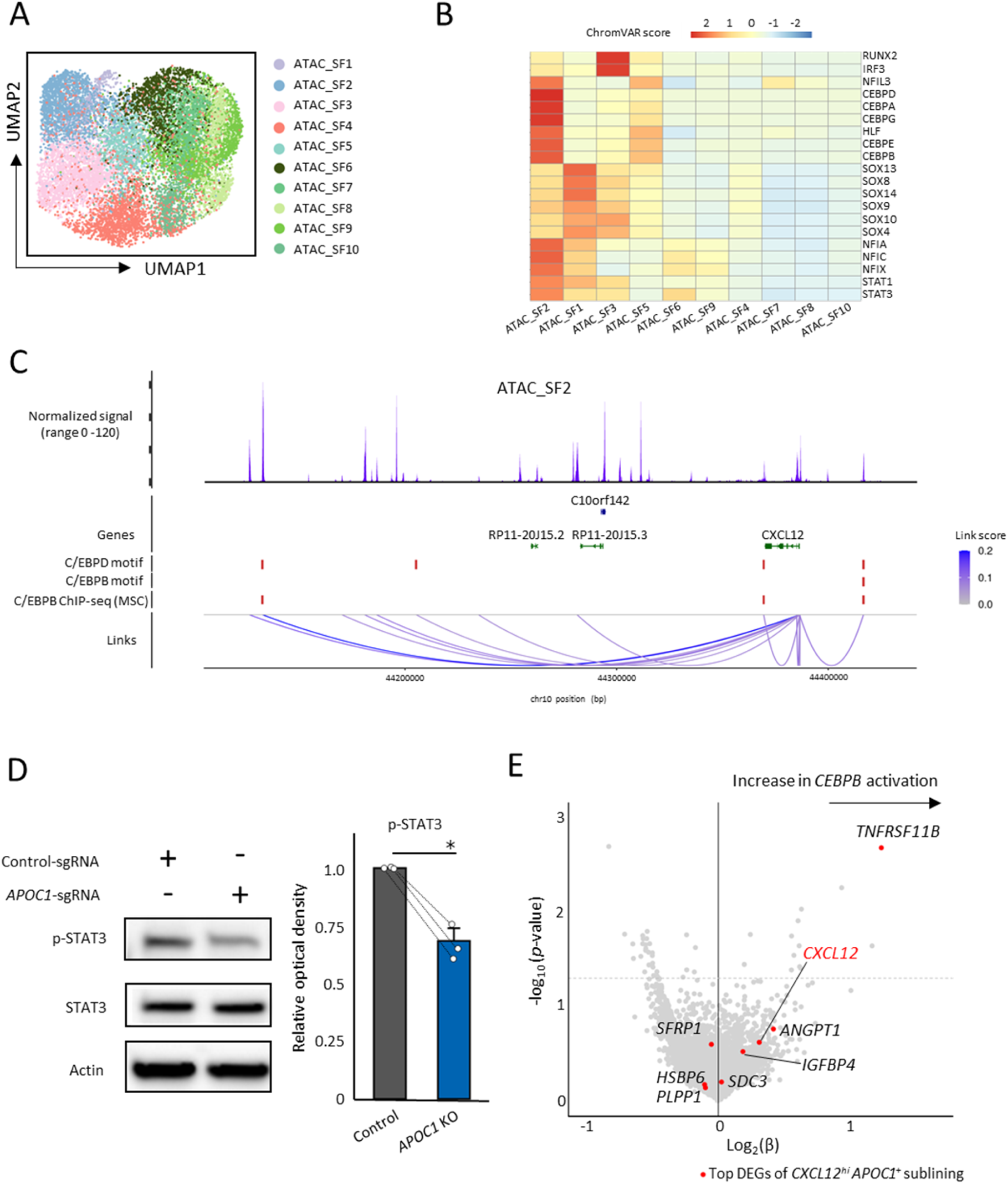
Multi-omics analyses identified the STAT3–C/EBP axis as a core transcriptional program of *CXCL12*^hi^ *APOC1*^+^ fibroblasts. **(A)** Uniform manifold approximation and projection (UMAP) visualization of synovial fibroblast (SF) fine clusters defined by chromatin accessibility. **(B)** Chromatin accessibility, measured by chromVAR scores, of the top 20 transcription factors in ATAC_SF2 across SF clusters defined by single-nucleus assay for transposase-accessible chromatin sequencing (snATAC-seq). **(C)** Organization of transcriptional regulatory regions surrounding *CXCL12* in ATAC_SF2. Link color intensity represents the link score (correlation coefficient between gene expression and accessibility of peaks located within ±500 kb of the transcription starting site (TSS)). Highlighted peaks include those containing *C/EBPB* and *C/EBPD* motifs (link score > 0.05, *p* < 0.05) and overlapping with public C/EBPβ ChIP-seq peaks in mesenchymal stromal cells (SRX1027438, SRX1027439). *P* values correspond to the z-score-based significance of correlation coefficients relative to background peaks. **(D)** Expression of phosphorylated STAT3 (phospho-STAT3) and total STAT3 in SF depleted of *APOC1* (blue) or negative control (black). Protein levels were analyzed by western blotting, and phospho-STAT3 was quantified relative to total STAT3 and normalized to control samples. Bars, mean; error bars, SEM. *P* values, a paired t-test from three independent experiments (* *p* <0.05). **(E)** Volcano plot showing transcriptional changes in fibroblast Perturb-seq upon *CEBPB* overexpression, with the top 30 DEGs of *CXCL12*^hi^ *APOC1*^+^ sublining SF highlighted in red. ChIP-seq, chromatin immunoprecipitation sequencing; chr, chromosome; DEG, differentially expressed gene; kbp, kilobase pair; KO, knock out; MSC, mesenchymal stromal cell; RA, rheumatoid arthritis; SD, standard deviation; SEM, standard error of the mean; SF, synovial fibroblast; sgRNA, single-guide RNA; snATAC-seq, single-nucleus assay for transposase-accessible chromatin sequencing; TFs, transcription factors; TSS, transcription start site; UMAP, Uniform Manifold Approximation and Projection.

### *CXCL12*^hi^ *APOC1*^+^ sublining SF displays senescent fibroblast-like phenotype

We identified the STAT3-C/EBP axis as a core transcriptional regulator of *CXCL12*^hi^ *APOC1*^+^ sublining SF. This is notable because the C/EBP family have recently emerged as central mediators of SASP(*41*), and *CXCL12*^hi^ *APOC1*^+^ sublining SF expresses canonical SASP factors, including *IL6* and *CXCL12*(*41*), together with cognate signaling components such as *IL6R*, *IL6ST* and *ACKR3* (supplemental figure S12A, figure 3B). These findings suggest a local autocrine or paracrine inflammatory circuit that may persist independently of immune cells-derived cytokines. Secreted frizzled-related protein 1 (SFRP1), a secreted antagonist of Wnt signaling and a SASP factor induced by DNA damage or oxidative stress(*42*), is also specifically expressed by *CXCL12*^hi^ *APOC1*^+^ sublining SF (supplemental figure S4F). Moreover, CXCL12 is a recognized SASP component produced by senescent cancer cells and CAF, where it promotes tumor growth, angiogenesis and immune cell infiltration across multiple malignancies(*43*). In support of this link, Meguro *et al* reported that the p16^high^ senescent (p16^h^-sn) fibroblasts in the aged mouse bladder establish an iCAF-like, tumor-permissive niche through CXCL12 secretion(*44*). GSEA revealed that genes upregulated in p16^h^-sn fibroblasts(*44*) (supplemental table 9) were significantly enriched among DEGs suppressed by *APOC1* knockdown in RA-SF (enrichment score = 0.358, adjusted *p* < 0.001; figure 6, A and B), suggesting that APOC1 promotes a senescent CAF-like program in RA-SF. Consistently, our scRNA-seq data showed that *CXCL12*^hi^ *APOC1*^+^ sublining SF highly express a gene signature of p16^h^-sn CAFs associated with aging and poor prognosis in bladder cancer(*44*) (supplemental figure S12B). Spatial analysis further demonstrated that *CXCL12*^hi^ *APOC1*^+^ sublining SF forming plasmablast-rich niches partially expressed *CDKN1A* and *CDKN2A*, established markers of senescence (supplemental figure S12C). Collectively, these findings indicate that cellular senescence represents a key feature of *CXCL12*^hi^ *APOC1*^+^ sublining SF.

**figure 6.**
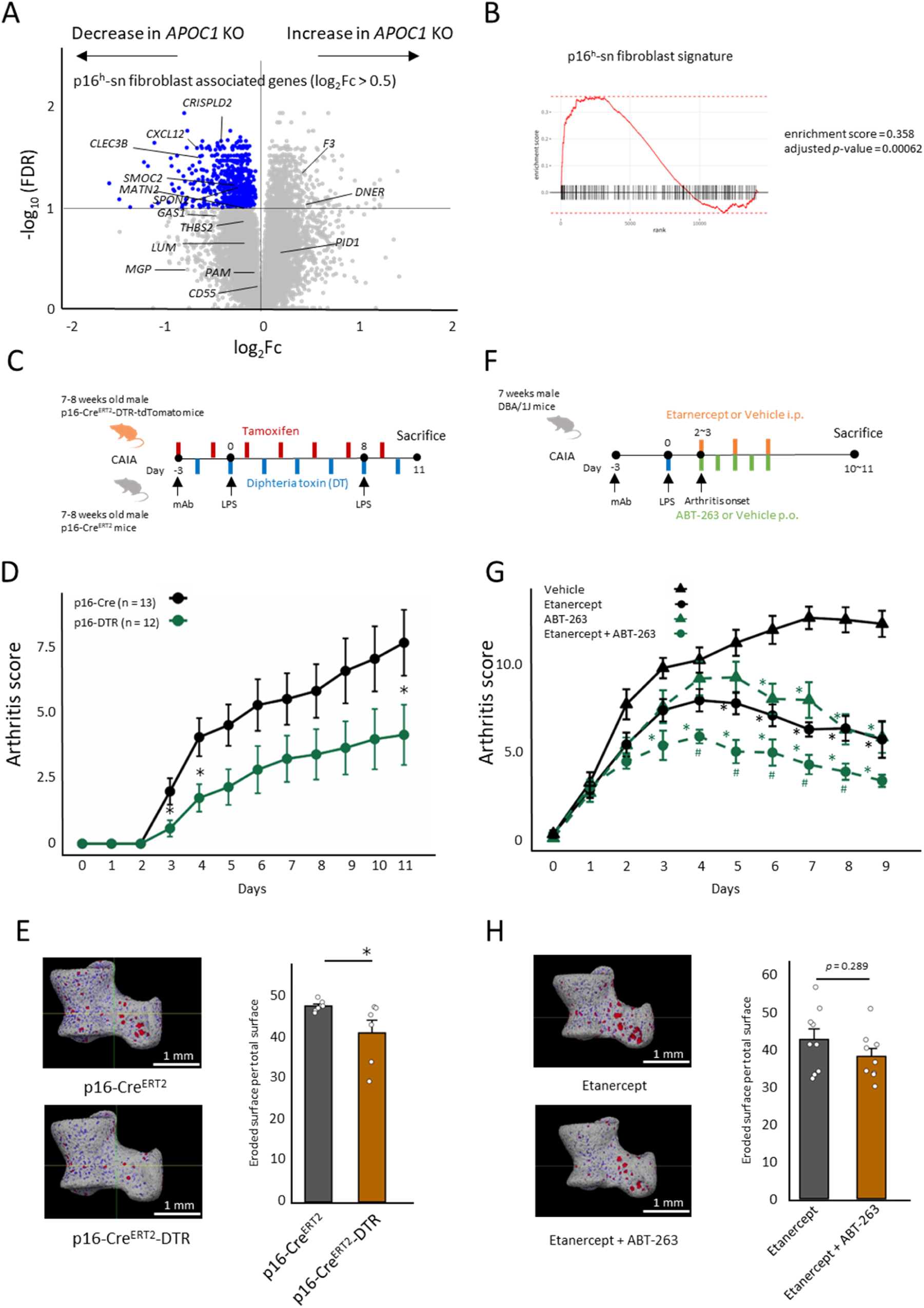
Elimination of senescent cells contributes to the alleviation of experimental synovitis. **(A)** Differential gene expression analysis comparing synovial fibroblasts (SF) depleted of *APOC1* or negative control. Blue points, genes with significantly decreased expression (false discovery rate (FDR) < 0.01); black labels, marker genes of p16^h^-sn fibroblast from Meguro *et al*(*44*). **(B)** Enrichment of p16^h^-sn fibroblast marker genes in negative control relative to *APOC1*-depleted SF, calculated by Gene Set Enrichment Analysis (GSEA). **(C)** Experimental design of the collagen antibody-induced arthritis (CAIA) mouse model using 7-8 week-old male p16-Cre^ERT2^-DTR mice or p16-Cre^ERT2^ mice (control). Starting at the time of arthritis induction, mice received intraperitoneal injections of tamoxifen 80 mg/kg and diphtheria toxin 25 μg/kg twice daily. **(D)** Clinical arthritis scores of p16-Cre^ERT2^-DTR mice (n = 12) and p16-Cre^ERT2^ control mice (n = 13) under CAIA condition. Dots, mean; error bars, SD. *P* values, two-tailed Mann-Whitney *U* test (* *p* <0.05). **(E)** Ratio of eroded surface to total surface area of left ankle joints in p16-Cre^ERT2^-DTR mice (n = 12) and p16-Cre^ERT2^ control mice (n = 13) under CAIA condition, assessed by micro-computed tomography (micro-CT). Bars, mean; error bars, SD. *P* values, two-tailed Mann-Whitney *U* test (* *p* <0.05). **(F)** Experimental design of the CAIA mouse model using 7-week-old male DBA/1J wild-type mice treated with the TNF inhibitor etanercept and/or the senolytic agent ABT-263. Following arthritis onset, mice received intraperitoneal injections of vehicle or etanercept 2 mg/kg and oral administration of vehicle or ABT-263 100 mg/kg once daily. **(G)** Clinical arthritis scores of mice treated with etanercept plus ABT-263 (n = 9), etanercept alone (n = 9), ABT-263 alone (n = 8) or vehicle (n = 9) under CAIA condition. Dots, mean; error bars, SD. *P* values, two-tailed Mann-Whitney *U* test (* *p* < 0.01 versus vehicle; ^#^ *p* < 0.05 for etanercept plus ABT-263 versus etanercept alone). **(H)** Ratio of eroded surface to total surface area of left ankle joints in mice treated with etanercept plus ABT-263 (n = 9) or etanercept alone (n = 9) under CAIA condition, assessed by micro-CT. Bars, mean; error bars, SD. *P* values, two-tailed Mann-Whitney *U* test (* *p* <0.05). CAIA, collagen antibody-induced arthritis; CT, computed tomography; DT, diphtheria toxin; DTR, diphtheria toxin receptor; FDR, false discovery rate; GSEA, Gene Set Enrichment Analysis; ip, intraperitoneal; iv, intravenous; KO, knock out; LPS, lipopolysaccharide; po, per os; SD, standard deviation; SF, synovial fibroblast; sn, senescent; TNF, tumor necrosis factor.

### Prophylactic and therapeutic elimination of senescent cells alleviates experimental arthritis

We next examined whether senescent cells contribute to synovitis progression *in vivo*. Collagen antibody–induced arthritis (CAIA) was established in p16-Cre^ERT^-tdTomato mice(*44*) to trace p16^high^ (p16^h^) cells in inflamed synovium following tamoxifen (TAM) administration (figure 1E; supplemental figure S12D).

Tomato^+^ synovial cells co-expressing podoplanin (PDPN) were detected in the sublining layer (supplemental figure S12E). Whereas Tomato^+^ cells constitute well below 1% of total cells in normal tissues of young mice(*45*), FCM analysis revealed that Tomato^+^ SF were expanded to approximately 1% of total SF in arthritic joints.

To determine the functional contribution of p16^h^ cells, CAIA was induced in p16-Cre^ERT2^-DTR mice, with p16-Cre^ERT2^ littermates as controls (figure 6C). Selective ablation of p16^h^ cells by TAM and diphtheria toxin (DT) administration(*45*) significantly attenuated arthritis severity (figure 6D) and reduced cartilage destruction and bone erosion, as assessed by micro-computed tomography (micro-CT) and histopathological analysis (figure 6E and supplemental figure S12, F to I).

We next evaluated pharmacological senolysis using navitoclax (ABT-263), a BCL-2 family inhibitor with potent senolytic activity in stromal cells(*46*) and efficacy in refractory myeloproliferative neoplasms in clinical trials(*47*). Notably, *APOC1* knock-down reduced *BCL2* expression (supplemental figure S13A)—a key target of ABT-263—while activating cell-cycle–associated programs (supplemental figure S13, A and B), suggesting that APOC1 supports a senescent fibroblast state linked to cell-cycle arrest with BCL-2–mediated apoptotic resistance(*48*). In wild-type DBA/1J mice with established CAIA, ABT-263 monotherapy significantly ameliorated arthritis, and combined ETN and ABT-263 treatment provided greater benefit than ETN alone (figure 6, F to H; supplemental figure S13, C to F). These findings indicate that senolytic intervention provides additive therapeutic benefit when combined with cytokine blockade *in vivo*.

Collectively, these data demonstrate that senescent iCAF-like SF contribute to arthritis pathogenesis, highlighting senolytic targeting as a potential therapeutic strategy for treatment-refractory inflammatory arthritis.

## Discussion

Our multi-omics analysis across 54 RA patients identifies a previously unrecognized fibroblast state in refractory RA synovium, refining the current framework of SF heterogeneity. Although SF highly expressing CXCL12 and IL-6 have conventionally been categorized as inflammatory fibroblasts(*10*), our study reveals that *CXCL12*^high^ fibroblasts are not a homogeneous population. Instead, we demonstrate that a distinct *APOC1*-expressing subset defines a unique iCAF-like program associated with treatment resistance.

*CXCL12*^hi^ *APOC1*^+^ sublining SF represents distinct transcriptional, spatial, and functional features, and is enriched in patients with poor response to cytokine-blocking agents, and forms a niche with plasmablasts via the CXCL12–CXCR4 axis, suggesting a central role in sustaining refractory synovitis. Importantly, *CXCL12*^hi^ *APOC1*^+^ sublining SF shows little enrichment of canonical cytokine-responsive programs and is largely refractory to modulation by current disease-modifying therapies, suggesting that this cluster cannot be fully explained by conventional inflammation-driven paradigms. From a mechanistic perspective, our data support the notion that this iCAF-like state is governed by a STAT3–C/EBP transcriptional cascade and is closely linked to active SASP production, with APOC1 as both a defining marker and a modulator of this broader iCAF-like program. Although stromal senescence remains poorly characterized in inflammatory joint disease, prior studies have suggested the presence of senescent synovial cells in RA(*49*), and senolytic agents such as navitoclax (ABT-263) or UBX0101 have shown benefit in osteoarthritis models(*50*). Our findings extend these observations and support senescence-associated stromal programs as potential therapeutic targets in refractory RA.

This study has several limitations. First, the relatively small sample size and heterogeneous patient backgrounds limit the robustness of prognostic analyses within our cohort, although external validation using public bulk RNA-seq datasets partially mitigates this concern. Second, *in vivo* ablation of p16^hi^ cells cannot be attributed exclusively to senescent SF, as other p16-expressing cell populations may also contribute to the observed phenotype. Third, it has not been formally demonstrated that *CXCL12*^hi^ *APOC1*^+^ sublining SF uniquely account for the senescent phenotype within the broader SF compartment. Future studies should delineate the senescence-associated properties of this subset and determine whether its selective depletion enhances responsiveness to cytokine-blocking therapies.

In summary, our single-cell multi-omics approach reveals stromal cell senescence as a critical component of treatment-resistant RA pathophysiology (supplemental figure S14). Therapeutic strategies targeting iCAF-like fibroblasts with senescent properties, coupled with patient stratification based on synovial senescence signatures, may provide new avenues for patients refractory to existing cytokine-directed therapies.

## Methods Study

### design

Synovial tissue samples were obtained from 54 patients with RA fulfilling the 2010 ACR/EULAR and/or the 1987 American College of Rheumatology classification criteria. Flow cytometry, single-cell transcriptomics, single-nucleus multiome profiling, and spatial transcriptomics were integrated with prospective clinical data to identify fibroblast states and immune niches associated with treatment-refractory synovitis. Mechanistic studies using cultured RA-SF, targeted knockdown experiments, and *in vivo* arthritis models were performed to define molecular pathways underlying pathogenic fibroblast reprogramming and therapeutic resistance. Detailed experimental procedures are provided in the Supplementary Materials and Methods.

### Statistical analysis

Statistical analyses were performed using appropriate parametric or non-parametric tests, as indicated. For *in vitro* experiments, comparisons between two groups were conducted using paired or unpaired *t*-tests, Wilcoxon signed-rank tests, or Mann–Whitney U tests, as appropriate. Comparisons among more than two groups were performed using the Kruskal–Wallis test followed by Dunn’s multiple-comparison test with Bonferroni correction. A *p* value <0.05 was considered statistically significant unless otherwise stated. Bar graphs are presented as mean ± standard deviation (SD). For large-scale analyses, including differential gene expression analyses, multiple-testing correction was performed using the Benjamini–Hochberg method to control the false discovery rate.

## Acknowledgments

The super-computing resource was provided by Human Genome Center, Institute of Medical Sciences, The University of Tokyo (http://sc.hgc.jp/shirokane.html). We are deeply grateful to the patients who participated in this study. We thank the laboratory members including Mayuko Fukuda and Haruka Takahashi at Department of Allergy and Rheumatology, Graduate School of Medicine, The University of Tokyo, for assistance with experiments and data collection, and for helpful discussions. We appreciate the technical expertise provided by Junko Zenkoh and Kazumi Abe at Department of Computational Biology and Medical Sciences, Graduate School of Frontier Sciences, The University of Tokyo.

## Author contributions

Conceptualization: RYo, HT, KF; Data curation: RYo, RYa, IT, TY; Methodology: RYo, RYa, IT, YO, YS, TO, HT; Investigation: RYo, RYa, IT, SN, HT, YO, TMa, ST, YY; Formal analysis: RYo, RYa, TY, KI, TI, MO, KY; Visualization: RYo, TY; Resources: TWW, MN, YF, HS, KY, NT, TMi, KO, AM, TA, YT, and TT; Supervision: TWW, MN, YO, TMa, ST, KI, TI, MO, TO, HT, KF; Funding acquisition: RYo, HT, KF; Project administration: HT, TO, KF; Writing – original draft: RYo, HT; Writing – review & editing: RYo, IT, TY, TWW, MN, YO, ST, KI, TI, MO, KY, YY, YF, HS, NT, TMi, KO, AM, TA, YT, TT, YS, TO, HT, KF; Validation: RYo, SN, IT, HT.

## Data availability

The single-cell RNA-seq data generated for this study have been deposited in NBDC hum0307; JGAS000899. Public perturb-seq of fibroblast dataset from Southard *et al* was downloaded from https://doi.org/10.5281/zenodo.15200179.

## Code availability

The codes used for this article are available on Zenodo (https://doi.org/10.5281/zenodo.19491739).

