## Supplementary document for "An *APOC1*^+^ inflammatory CAF-like state drives a senescent, treatment-resistant niche in rheumatoid arthritis"

**Supplementary materials and methods**

**Collection and quality control of synovial tissue samples from patients with rheumatoid arthritis**

This study was approved by the Ethics Committees of the University of Tokyo (approval numbers G10137 and 2021057G) and the participating medical institutions. Written informed consent was obtained from all participants in accordance with the Declaration of Helsinki. Synovial tissue samples were obtained from 54 patients with rheumatoid arthritis (RA), yielding a total of 56 specimens. Of these, 44 samples were collected by ultrasound-guided synovial biopsy and 10 samples were obtained during arthroplasty surgery. The cohort included 16 treatment-naive patients, 35 patients receiving conventional synthetic disease-modifying antirheumatic drugs (csDMARDs), and 14 patients treated with biologic or targeted synthetic DMARDs (b/tsDMARDs) for more than three months (supplemental figure S1 and supplemental table 1). At the time of biopsy, 46 of the 54 patients exhibited moderate to high disease activity. Disease activity was further monitored for three months following biopsy and treatment modification.

All patients were aged ≥18 years and fulfilled the diagnostic criteria for RA according to the 2010 ACR/EULAR classification criteria and/or the 1987 American College of Rheumatology criteria. Ultrasound-guided synovial biopsies were performed under aseptic conditions by experienced orthopaedic surgeons using an 18-gauge Mission Core biopsy needle (BD, 227ADBZX00049000). Joints targeted for biopsy had not received intra-articular injections within four weeks before the procedure or during sampling. No procedure-related adverse events were observed. At least three synovial tissue fragments from each sample were formalin-fixed and paraffin-embedded. Remaining tissue was sectioned into approximately 1 $mm^3$ fragments and cryopreserved in CryoStor CS10 (STEMCELL Technologies, 100-1061) using a Mr. Frosty freezing container (Thermo Fisher Scientific, 5100-0001). To assess sample quality, haematoxylin and eosin (H&E)–stained sections were independently reviewed by experienced pathologists. Samples were excluded if they lacked an identifiable synovial lining layer or consisted solely of loose fibrovascular tissue without a lining structure.

**Categorization of synovial pathotypes by immunohistochemistry**

For downstream analyses, synovial samples were classified into three distinct pathotypes based on immunohistochemical (IHC) staining, as previously described by Humby *et al*(*1*). Synovial tissue sections were stained with antibodies against CD3 (clone LN10; Leica Biosystems), CD20 (clone L26; Leica Biosystems), CD138 (clone B-A38; Nichirei Biosciences), and CD68 (clone PG-M1; Agilent). Pathotype assignment was performed according to the relative abundance and spatial distribution of T cells, B cells, plasma cells, and macrophages within the synovial tissue.

**Enzymatic dissociation, flow cytometry and single-cell RNA-sequencing (scRNA-seq) library preparation of RA synovium**

Cryopreserved synovial tissue samples were thawed using ThawSTAR (Biolife Solutions) and enzymatically dissociated in Advanced Dulbecco's modified Eagle's medium (DMEM; Gibco, 12491015) supplemented with Liberase TL (0.1 mg/ml; Roche, 5401020001) and DNase I (0.1 mg/ml; Roche, 10104159001) for 15 min at 37 °C in a water bath. Following digestion, single-cell suspensions were generated by gentle mechanical dissociation using a cell scraper and filtration through a 70 μm cell strainer. Cells were subjected to Fc receptor blocking using Human TruStain FcX (BioLegend, 422302) and subsequently stained with (i) flow cytometry antibodies (supplemental table 10), (ii) oligonucleotide-barcoded antibodies for Cellular Indexing of Transcriptomes and Epitopes by Sequencing (CITE-seq), including cell hashing antibodies (supplemental table 11), and (iii) LIVE/DEAD Fixable Aqua Dead Cell Stain (Invitrogen, L34957). Samples were analyzed and sorted by fluorescence-activated cell sorting (FACS) on a BD FACSAria Fusion (BD). Up to 10,000 live $CD45^+$ cells ($CD45^+$ $Aqua^-$) and 10,000 live non–red blood cells from the $CD45^-$ fraction ($CD45^-$ $CD235a^-$ $Aqua^-$) were collected per sample. The gating strategy is shown in supplemental figure S2. Flow cytometry data were analyzed using FlowJo v10.10.0. Sorted live cells were loaded onto a Chromium Next GEM Chip K (10x Genomics, PN-1000287) and processed using the *Chromium Single Cell 5' Reagent Kits v2 (dual index)* according to the manufacturer's instructions (10x Genomics, CG30031). Gene expression (GEX) libraries were sequenced to a depth of at least 30,000 read pairs per cell, and antibody-derived tag (ADT) libraries were sequenced to at least 5,500 read pairs per cell. Sequencing was performed on a NovaSeq 6000 platform (Illumina) using 150-bp paired-end reads.

**scRNA-seq data analysis**

Raw sequencing data (FASTQ files) were processed using Cell Ranger multi v6.1.2 (10x Genomics). Briefly, reads were demultiplexed, and individual cells were assigned to their respective donors based on sample barcodes and cell hashing information. Reads were aligned to the Genome Reference Consortium Human Reference 38 (GRCh38), and unique molecular identifier (UMI) count matrices were generated for both GEX and ADT libraries. Putative doublets were identified and removed using scDblFinder(*2*). Downstream analyses were performed using Seurat v3(*3*). Quality control filtering excluded cells with mitochondrial gene content exceeding 5% or fewer than 500 detected genes. After filtering, a total of 82,708 cells were retained for subsequent analyses. Gene expression data were normalized using SCTransform and the 3,000 most variable features were identified.. Data integration across 53 datasets was performed by identifying integration anchors using the FindIntegrationAnchors() function, followed by data integration with IntegrateData() to generate a unified expression matrix. Principal component analysis (PCA) was conducted on the integrated dataset, and the first 25 principal components were used for downstream clustering and dimensionality reduction. A shared nearest neighbor (SNN) graph was constructed using the FindNeighbors() function in Seurat v3(*3*), and clusters were identified using the FindClusters() function with a resolution parameter of 0.7. This analysis yielded 29 distinct clusters, including clusters corresponding to low-quality cells and doublets, which were visualized using uniform manifold approximation and projection (UMAP). Cell type annotation was performed manually based on established marker genes and prior literature. Differentially expressed genes (DEGs) for each cluster were identified using the FindAllMarkers() function. For subclustering of synovial fibroblast (SF), only SF clusters were extracted, and PCA and UMAP analyses were repeated using 18 principal components with a clustering resolution of 0.4. Gene set activity scores were calculated at the single-cell level using the AddModuleScore() function implemented in Seurat.

**Definition of Multi-Omics and MOdular Stromal And Immune Cell (MOSAIC) scores**

The relative abundance of key synovial cell populations, including $CD3^+$ T cells, $CD19^+$ B cells, monocytes, SF, and endothelial cells, was quantified as the proportion of each cell type among total live cells. Based on scRNA-seq-derived cell clusters, we calculated (i) the proportion of major hematopoietic clusters relative to total hematopoietic cells, (ii) the proportion of major non-SF stromal clusters relative to total non-SF stromal cells, and (iii) the proportion of fine SF clusters relative to total SF. To identify co-varying cell populations, hierarchical clustering was performed on these proportional data using Euclidean distance and Ward's D2 linkage method, resulting in seven distinct cell modules. For each cell module, multidimensional scaling (MDS) was applied, and the first dimension was defined as the module score. To ensure consistency in the directionality of the scores, module scores were multiplied by −1 when the correlation coefficient between the module score and the corresponding cell-type proportion was negative.

**Gene set variation analysis applied to synovial fibroblast fine clusters**

To characterize cytokine responsiveness and drug-associated transcriptional programs in SF fine clusters, we leveraged publicly available bulk RNA-seq datasets of cultured RA-SF stimulated with individual cytokines(*4*) or treated with disease-modifying agents(*5*). For each cytokine condition, the top 1,000 significantly upregulated genes were defined as cytokine-stimulation signature genes (supplemental table 2). Similarly, for each drug treatment, the top 1,000 significantly downregulated genes were defined as medication-suppressive signature genes (supplemental table 3). We also extracted top-ranked genes of each cancer-associated fibroblast (CAF) cluster from DEG lists of published CAF scRNA-seq studies(*6-10*) (supplemental table 7). Pseudo-bulk gene expression profiles for each SF fine cluster were generated by aggregating single-cell expression data using the PseudoBulkExpression() function implemented in Seurat v5(*11*). Gene set variation analysis (GSVA) was then performed on the pseudo-bulk expression matrices to calculate enrichment scores for cytokine-stimulation signatures, medication-suppressive signatures, and top DEGs from CAF clusters(*6-10*) in each SF fine cluster.

**Receiver operating characteristic (ROC) analysis for treatment response prediction**

ROC analysis was performed using the pROC package in R(*12*) to assess the discriminatory performance of the proportion of *CXCL12*$^{hi}$ *APOC1*$^{+}$ fibroblasts among all SFs for CDAI50 response among patients treated with IL-6 receptor or TNF inhibitors. The analysis was restricted to 22 patients whose scRNA-seq datasets included 30 or more cells annotated as SF. CDAI50 response was defined as ≥50% improvement in CDAI from baseline to 3 months. AUCs and 95% confidence intervals were calculated using DeLong's method, and the optimal cutoff was determined by the Youden index.

**Mixed-effects association analysis of single-cell data to evaluate treatment response and synovial fibroblast fine clusters**

To assess the association between SF fine clusters and treatment response, we applied mixed-effects association of single cells (MASC)(*13*) to the scRNA-seq data. The analysis was restricted to 25 patients whose scRNA-seq datasets contained one or more cells annotated as SF. Associations were evaluated between the abundance of each SF fine cluster and clinical disease activity, as measured by the Clinical Disease Activity Index (CDAI), three months after initiation of interleukin-6 receptor (IL-6R) inhibitor or tumor necrosis factor (TNF) inhibitor therapy. For each SF fine cluster, a logistic regression model was fitted in which cluster membership of individual cells was treated as the outcome variable. Sex, age, baseline CDAI, percentage of mitochondrial gene expression, and total UMI counts were included as fixed-effect covariates. Donor identity and library preparation batch were incorporated as random-effect covariates to account for inter-individual variability and technical effects.

**Validation analysis using public datasets from the R4RA and STRAP trials**

For validation analyses, we extracted the top 15 marker genes for each SF fine cluster identified using the FindAllMarkers() function in Seurat v5(*11*). Publicly available transcriptomic and clinical data were obtained from the R4RA trial (https://r4ra.hpc.qmul.ac.uk/) and the STRAP trial cohorts (https://strap.hpc.qmul.ac.uk/). For each gene, Spearman's rank correlation coefficients between baseline gene expression levels and changes in the CDAI at 16 weeks after treatment initiation were extracted (supplemental table 4 and 5). To assess whether SF fine cluster–specific gene signatures were systematically associated with treatment response, a binomial test was performed to determine whether the distribution of correlation coefficients for the 15 genes within each cluster showed a significant bias toward positive or negative values for each therapeutic agent. Multiple hypothesis testing was controlled using the Bonferroni correction.

**Single-cell multiome library preparation of RA synovium**

Synovial tissue samples from nine patients with RA were cryopreserved and subsequently thawed, enzymatically digested, stained, and fluorescence-activated cell sorted as described above for scRNA-seq analyses. Following cell sorting, nuclei were isolated using a modified version of the *Nuclei Isolation for Single Cell Multiome ATAC + Gene Expression Sequencing protocol* (CG000365 Rev. C; 10x Genomics). Approximately 10,000 nuclei per sample were incubated with the transposition reaction mix and subsequently loaded onto a Chromium Next GEM Chip J, where they were partitioned into Gel Beads-in-Emulsion (GEMs). GEM incubation and library construction were performed according to the *Chromium Next GEM Single Cell Multiome ATAC + Gene Expression protocol* (CG000338 Rev. F; 10x Genomics). Assay for transposase-accessible chromatin (ATAC) libraries were sequenced to a depth of at least 40,000 read pairs per nucleus, and gene expression libraries were sequenced to a depth of at least 30,000 read pairs per nucleus using a NovaSeq X Plus platform (Illumina).

**Data processing, quality control, batch correction, clustering and visualization of the multiome dataset**

Raw FASTQ files were initially processed using Cell Ranger ATAC v2.0.2 (10x Genomics), which performs barcode processing and read alignment to the human reference genome (hg38). Five of the nine samples were multiplexed and subsequently demultiplexed into individual donors using demuxlet(*14*) based on single-nucleotide polymorphisms (SNPs) genotyped with the Japanese Screening Array (Illumina). Downstream analyses were performed using Seurat v5(*11*) and Signac v1.14.0(*15*) to generate Seurat objects containing paired gene expression and chromatin accessibility profiles for each nucleus. Quality control filtering retained nuclei with 500–50,000 ATAC fragments, 500–25,000 detected transcripts, nucleosome signal <2, and transcription start site (TSS) enrichment >4. After filtering, a total of 26,430 single nuclei were retained for further analyses. For the RNA modality, gene expression counts were normalized and variance-stabilized using SCTransform, followed by principal component analysis (PCA). Batch effects across samples were corrected using reciprocal PCA (RPCA)-based integration. For the ATAC modality, peak-by-cell count matrices were normalized using term frequency–inverse document frequency (TF–IDF) transformation implemented in

Signac. Peaks with low average accessibility were removed using the FindTopFeatures() function prior to dimensionality reduction. Singular value decomposition (SVD) was applied to the TF-IDF matrix using the RunSVD() function to generate a latent semantic indexing (LSI) representation. To correct batch effects in chromatin accessibility data, integration anchors derived from the RNA modality were reused to integrate the LSI embeddings. LSI dimensions 2-30 were used to construct a nearest-neighbor graph and to perform clustering. SF clusters were subsequently subsetted from the integrated object and re-clustered using the same LSI dimensions at a resolution of 0.6. ATAC-based clusters were annotated based on the expression of canonical marker genes from the RNA modality. TF motif annotations were added using the JASPAR2022(*16*) database and the hg38 reference genome. Motif accessibility deviations were quantified using the RunChromVAR() function after restricting peaks to chromosomes present in the hg38 BS genome. GC content and other base composition statistics were computed using the RegionStats() function in Signac. Peak-to-gene links were inferred using the LinkPeaks() function to associate chromatin accessibility with gene expression, and links were filtered using a threshold of $P < 0.05$ and a link score $>0.05$. TF motif occurrences within linked peaks were identified based on JASPAR2022 annotations.

**Reference mapping using Symphony**

To assess the concordance our multiome dataset of with our scRNA-seq dataset, reference-based mapping was performed using Symphony(*17*). A Symphony reference was constructed from our scRNA-seq dataset of SF using the buildReference() function. Query SF derived from the multiome dataset were subsequently mapped onto this internally generated reference using the mapQuery() function implemented in Symphony, and reference cell types and transcriptional states were predicted for each query cell using the knnPredict() function with $k = 30$.

**Xenium sample preparation and Xenium analyzer processing**

Formalin-fixed, paraffin-embedded (FFPE) tissue sections were mounted onto Xenium slides and processed according to the *Xenium In Situ for FFPE Tissue Preparation Guide* (CG000578 Rev. C, 10x Genomics). Deparaffinization and decrosslinking were performed following the *Xenium In Situ for FFPE*

*Deparaffinization and Decrosslinking protocol* (CG000580 Rev. D; 10x Genomics). Briefly, slides were baked at 60 °C for 30 min, followed by sequential deparaffinization in xylene, rehydration through graded ethanol, and rinsing in nuclease-free water. Tissue sections were then incubated in decrosslinking and permeabilization solution at 80 °C for 30 min and washed with PBS containing Tween-20 (PBS-T). Subsequent processing was carried out according to the *Xenium Prime In Situ Gene Expression protocol* (CG000760 Rev. C; 10x Genomics). Probes from the Xenium Prime 5K Human Pan-Tissue panel, together with custom add-on panels (supplemental table 12), were hybridized to the tissue sections at 50 °C for 18 h. Following hybridization, slides underwent sequential washing, ligation, and rolling-circle amplification. Background fluorescence was chemically quenched, and slides were loaded into the Xenium Analyzer for imaging and data acquisition in accordance with the *Xenium Analyzer User Guide* (CG000584 Rev. G, 10x Genomics). After completion of Xenium imaging, slides were subjected to post-run H&E staining according to the *Xenium In Situ Gene Expression Post–Xenium Analyzer H&E Staining protocol* (CG000613 Rev. B; 10x Genomics). Following H&E staining, the same tissue sections were further processed for PhenoCycler-Fusion analysis.

**Cell segmentation, quality control and cell type annotation of Xenium data**

Cell segmentation of Xenium in situ gene expression data was performed using Xenium Onboard Analysis (10x Genomics). Segmentation was conducted using the manufacturer-recommended multimodal strategy, incorporating boundary marker stains (ATP1A1, CD45 and E-cadherin), an interior RNA marker (18S rRNA), and nuclear expansion where appropriate, as described in the *Xenium In Situ Multimodal Cell Segmentation: Workflow and Data Highlights* documentation. For quality control, segmented cells with ≤50 detected transcript species (nFeatures) were excluded from further analyses. The filtered Xenium dataset was subsequently imported into Seurat v5(*11*) for downstream analyses. Cell type annotation was performed by reference-based deconvolution using robust cell type decomposition (RCTD)(*18*). Our independently processed and annotated scRNA-seq dataset was used as the reference. Cell identities in the Xenium data were inferred based on transcriptional similarity to the reference profiles, restricted to the 5,101 genes included in the Xenium Prime 5K panel and the custom add-on panel. Xenium Explorer v3.2 was used for visualization and spatial exploration of selected cell types and molecular features.

**Neighborhood enrichment analysis and inference of cell–cell interactions**

Spatial neighborhood enrichment analysis was performed using the squidpy.gr.nhood_enrichment() function implemented in Squidpy v1.6.5(*19*) to assess whether specific cell types were preferentially localized in proximity to one another beyond random expectation. Statistical significance of cell-type co-localization was evaluated based on permutation testing as implemented in the package. Cell–cell communication analysis was conducted using CellChat v2.1.2(*20*). To incorporate spatial information, ligand–receptor interactions were inferred only between cells located within a radial distance of 250 µm. Communication probabilities were computed using the computeCommunProb() function, thereby restricting interaction inference to spatially proximal cell pairs.

**Multiplexed immunostaining by PhenoCycler-Fusion**

Following Xenium analysis, the same tissue sections were subsequently processed for multiplexed immunostaining using the PhenoCycler-Fusion system (Quanterix) according to the manufacturer's instructions. Antigen retrieval was performed by immersing slides in Tris–EDTA buffer (pH 9.0) and heating in a pressure cooker for 20 min. Tissue sections were then incubated with a panel of 60 antibodies for 3 h at room temperature (supplemental table 13). After staining, sections were washed and antibody binding was chemically fixed. Reporter cycling reactions and image acquisition were carried out using the PhenoCycler-Fusion instrument (Quanterix). Fluorescence images were acquired using a BZ-X810 fluorescence microscope (Keyence).

**Assessment of correlation between cell density and protein distribution**

To quantify the spatial enrichment of each cell type, Gaussian kernel density estimation (KDE) was performed using the KernelDensity() function implemented in scikit-learn v 1.7.2 (https://jmlr.csail.mit.edu/papers/v12/pedregosa11a.html). KDE maps were generated on a two-dimensional spatial grid to estimate local cell density across each tissue section. In parallel, fluorescence intensities from PhenoCycler-Fusion images were averaged within the same spatial grid units used for KDE map generation.

To assess the relationship between cell density and protein distribution, Pearson's correlation coefficients were calculated between the KDE-derived cell density maps and the corresponding grid-averaged fluorescence intensity values.

**Cell culture**

The human RA-SF cell line MH7A was obtained from the RIKEN BioResource Research Center. Primary RA-SF were either purchased from a commercial supplier (Articular Engineering) or isolated from synovial tissue obtained during arthroplasty surgery and were used for *APOC1* knockdown experiments. For primary cell isolation, freshly collected synovial tissues were minced and enzymatically digested in Advanced DMEM (Gibco, 12491015) supplemented with Liberase TL (0.1 mg/ml; Roche, 5401020001) and DNase I (0.1 mg/ml; Roche, 10104159001) for 30 min at 37 °C in a humidified atmosphere containing 5% $CO_2$. Isolated SF were cultured in DMEM with L-glutamine and phenol red (Fuji-film, 044-29765) supplemented with 10% FCS (BioWest, S1820-500), 100 U/ml penicillin, and 100 µg/ml streptomycin (Gibco, 10378-016). Culture medium was replaced every three days, and cells were passaged upon reaching approximately 90% confluence. SF between passages 3 and 6 were used for electroporation, RNA sequencing, and western blot analyses.

**Knockdown of *APOC1* by CRISPR-Cas9-mediated genome editing**

*APOC1* knockdown in MH7A cells or primary RA-SF was performed using the CRISPR-Cas9 genome-editing system. Recombinant Cas9 protein (0.03 nmol; Invitrogen, A36498) and synthetic single-guide RNA (sgRNA; 0.12 nmol; Horizon Discovery) were introduced into MH7A cells (50,000 cells) or primary RA-SF (25,000 cells) by electroporation using a 4D-Nucleofector system (Lonza, V4XC-2032). A non-targeting control sgRNA (U-009501-01) and an sgRNA targeting *APOC1* (target sequence: GAAGGGTTCTAACCGCATCT; protospacer-adjacent motif (PAM): TGG; SG-011574-01) were obtained from Horizon Discovery. Genome-editing efficiency was assessed using Inference of CRISPR Edits (ICE) analysis and was consistently greater than 90% across experiments.

**Cell viability assay**

MH7A cells subjected to Cas9 protein and sgRNA transfection were seeded into 96-well plates at a density of 2,000 cells per well. Cell viability was assessed over a 72-h period using the Cell Counting Kit-8 (CCK-8, Dojindo. CK04) according to the manufacturer's instructions. Briefly, 10 µl of CCK-8 solution was added to each well, followed by incubation for 4 h at 37 °C. Absorbance at 450 nm was measured using a Model 680 Series microplate reader (Bio-Rad). The experiment was independently performed four times.

**Migration and invasion assays**

Cell migration and invasion were assessed using Transwell-based assays. Migration assays were performed using 8-µm pore size Transwell chambers (Corning, 3422), and invasion assays were conducted using BioCoat Matrigel invasion chambers (Corning, 354480). MH7A cells transfected with Cas9 protein and sgRNA were resuspended in serum-free medium, and 12,500 cells in 500 µl were seeded into the upper chambers of 24-well plates. The lower chambers were filled with complete culture medium as a chemoattractant. Cells were allowed to migrate or invade for 48 h at 37 °C, after which non-migrated cells were removed and cells on the lower surface of the membranes were fixed and stained using a Diff-Quik staining kit (Sysmex, 16920). Migrated or invaded cells were quantified by counting cells in three randomly selected fields per membrane under ×400 magnification. Invasion efficiency was calculated as the percentage of invaded cells relative to migrated cells. All experiments were performed independently three times.

**Western blotting**

Western blotting was performed as previously reported(*21*) with minor modifications. Cultured human SF were lysed in buffer containing 1 % NP-40 (Milllipore, 492016), 50 mM Tris-HCl (pH 7.4, Millipore, T2194), and 150 mM NaCl (Promega, V4221) supplemented with protease and phosphatase inhibitors (Thermo Fisher Scientific, 78440). Protein concentrations were determined using the Pierce BCA Protein Assay Kit (Thermo Fisher Scientific, A65453) according to the manufacturer's instructions. Lysates were mixed with 2× Laemmli sample buffer (Bio-Rad, 1610737) and denatured by heating at 95 °C for 5 min. Equal amounts of protein (10 µg) were separated by SDS-PAGE on 4-15% gradient precast gels (Bio-Rad, 4561086) using Tris-glycine-SDS running buffer (Nippon Gene, 312-90321) and subsequently transferred onto PVDF membranes (Millipore,

IPVH10100). Membranes were blocked with BSA-based blocking solution (Thermo Fisher Scientific, 37520) and incubated overnight at 4 °C with primary antibodies diluted in TBS containing 3% BSA. The following primary antibodies were used at a dilution of 1:1,000: anti-STAT3 (clone: D3Z2G, Cell Signaling Technology, 9145), anti-phospho-STAT3 (clone: EP2147Y, Abcam, ab76315), and anti-actin (clone: 13E5, Cell Signaling Technology, 4970S). After washing four times with TBS containing 0.1% Tween-20 (TBS-T), membranes were incubated with HRP-conjugated anti-rabbit secondary antibody (concentration: 1:2000, Cell Signaling Technology, 7074S) diluted in TBS-T containing 5% skim milk (Fujifilm, 190-12865) for 45 min at room temperature. Membranes were then washed four additional times with TBS-T, and immunoreactive bands were detected using enhanced chemiluminescence (Cytiva, RPN2232). Band intensities were quantified using ImageJ(*22*).

**Bulk RNA sequencing of *APOC1*-knockdown synovial fibroblasts**

Primary RA-SF were subjected to *APOC1* knockdown by transfection and subsequently seeded at a density of 12,000 cells per well in 24-well plates. After 6 days of culture, cells were lysed in RLT buffer (Qiagen, 79216) supplemented with 2-mercaptoethanol (Sigma-Aldrich, M3148) and stored at -80 °C until further processing. Total RNA was extracted using the RNeasy Mini Kit (Qiagen, 74004) according to the manufacturer's instructions. Bulk RNA-seq libraries were prepared using the TruSeq Stranded mRNA Library Prep Kit (Illumina, 20020595). Libraries were sequenced on a NovaSeq X platform (Illumina) to generate 150-bp paired-end reads, yielding more than 60 million fragments per library.

**RNA-seq data processing and differential gene expression analysis of *APOC1*-knockdown synovial fibroblasts**

Sequencing adapters were removed from raw reads using Cutadapt (v 4.8; https://doi.org/10.14806/ej.17.1.200). Low-quality bases at the 3' ends (Phred quality score < 20) were trimmed using the FASTX-Toolkit (v0.0.14; http://hannonlab.cshl.edu/fastx_toolkit/), and reads containing more than 20% low-quality bases were discarded. Processed reads were aligned to the GRCh38 human reference genome using STAR (v2.5.3a)(*23*) in two-pass mode with gene annotations from GENCODE v27(*24*). For gene-level quantification, all transcript isoforms corresponding to the same gene were collapsed into a single feature as previously described(*25*). All samples achieved uniquely mapped read rates exceeding 90% and more than $1 \times 10^7$ uniquely mapped reads. Gene-level read counts were obtained using HTSeq (v2.0.7)(*26*). Lowly expressed genes were filtered using the filterByExpr() function, and library sizes were normalized across samples using the trimmed mean of M values (TMM) method implemented in edgeR(*27*). Normalized counts were log-transformed to counts per million (CPM) for downstream analyses. Differential gene expression analysis was performed using negative binomial generalized linear mixed models to evaluate the effect of *APOC1* knockdown. Donor identity was included as a random-effect covariate, and an offset term corresponding to the natural logarithm of the normalized library size was incorporated into the model. The normalized library size was calculated by multiplying the TMM normalization factors by the raw library sizes. Statistical significance of differential expression associated with *APOC1* knockdown was assessed using likelihood ratio tests comparing the full model with a null model. Genes with a false discovery rate (FDR) <0.05 were considered statistically significant.

Full Model:

$\text{Expr}_i \sim \text{Knock down}_i + (1|\text{Donor}_i) + \text{offset}(\log(\text{Normalization factor}_i \times \text{Library size}_i))$

Null Model:

$\text{Expr}_i \sim (1|\text{Donor}_i) + \text{offset}(\log(\text{Normalization factor}_i \times \text{Library size}_i))$

**Gene set enrichment analysis**

Gene set enrichment analysis (GSEA) was performed using fgsea v1.32.4(*28*). GSEA was applied to transcriptomic changes associated with *APOC1* knockdown in RA-SF. Genes were ranked based on the differential expression statistics derived from the negative binomial generalized linear mixed model described above. As a reference gene set, we used a curated gene signature upregulated in Tomato$^{+}$ (p16$^{h}$ sn) fibroblasts compared with Tomato$^{-}$ (p16$^{l}$) fibroblasts(*29*) (supplemental table 9) and top DEGs from iCAF in public scRNA-seq datasets of breast cancer(*8, 9*) (supplemental table 7), as reported previously. We defined cell-cycle–related pathways by extracting gene sets whose names contained “CELL_CYCLE”, “G2M”, “E2F”,

"MITOTIC" or "PROLIFERATION" from the Hallmark and Reactome collections of the Molecular Signatures Database (MSigDB)(*30*). Enrichment scores were calculated using permutation-based testing with 10,000 permutations. Statistical significance was assessed using nominal *p* values, followed by multiple-testing correction using the Benjamini–Hochberg procedure to obtain adjusted *p* values. Gene sets with an adjusted $p$ value < 0.05 were considered significantly enriched.

**Mice**

Mice were housed in groups of two to five per cage under controlled conditions (ambient temperature, 23–25 °C; humidity-controlled environment) with a 12-h light/dark cycle (lights on from 08:00 to 20:00). Animals were provided with standard chow (CA-1, CLEA), water ad libitum, and environmental enrichment. All procedures involving animals were conducted in accordance with the Guidelines for Animal Experiments of the Institute of Medical Science, the University of Tokyo (IMSUT), and were approved by the Animal Experiment Committee at IMSUT (approval numbers A16-33, A21-26 and A2025M025). All p16-Cre$^{ERT2}$-tdTomato or p16-Cre$^{ERT2}$-DTR mice were heterozygous and generated by crossing p16$^{INK4A}$-Cre$^{ERT2}$ mice(*31*) with Rosa26-CAG-LSL-tdTomato or Rosa26-SA-LSL-DTR-IRES-tdTomato reporter mice obtained from The Jackson Laboratory. For labeling of p16$^{h}$-sn cells in the p16-Cre$^{ERT2}$-tdTomato mouse model, tamoxifen (80 mg/kg body weight (BW), Cayman Chemical, 13258) dissolved in sunflower oil (Wako, 196-15265) was administered by intraperitoneal injection once daily for five consecutive days using a 25-gauge needle. Wild-type DBA/1J mice were purchased from Jackson Laboratory Japan.

**Arthritis induction**

Collagen antibody-induced arthritis (CAIA) was induced in p16-Cre$^{ERT2}$-tdTomato, p16-Cre$^{ERT2}$-DTR or p16-Cre$^{ERT2}$ mice (8-week-old male mice, background; C57BL/6) and wild type DBA/1J mice (7-week-old male mice). The number of animals used in each experiment is indicated in the corresponding figure legends. Briefly, mice were systemically immunized by i.p. injection of an anti-collagen monoclonal antibody cocktail (5 mg per mouse, Chondrex, 53040) on day 0. On day 3 and day 11, the p16-Cre$^{ERT2}$-tdTomato, p16-Cre$^{ERT2}$-DTR or p16-Cre$^{ERT2}$ mice received additional i.p. injections of lipopolysaccharide (LPS; 50 µg per mouse,

Chondrex, 53040). Wild type DBA/1J mice received LPS injections only on day 3. Mice were euthanized on day 14 and tissues were collected for downstream analyses as indicated.

**Treatments for senolysis**

For genetic senolysis, p16-Cre$^{ERT2}$-DTR or p16-Cre$^{ERT2}$ mice received alternating intraperitoneal (i.p.) injections of tamoxifen (80 mg/kg BW) and diphtheria toxin (DT; 25 µg/kg BW, Wako, 048-34371) for 14 consecutive days. Tamoxifen was dissolved in sunflower oil, and DT was diluted in phosphate-buffered saline (Nacalai Tesque, 14249-95). For pharmacological senolysis, wild-type DBA/1J mice were treated with etanercept (Pfizer, 4987123151559) or ABT-263 (Selleck, S1001) after the onset of collagen antibody-induced arthritis (CAIA) until the experimental endpoint. Etanercept was administered by i.p. injection at a dose of 2 mg/kg BW three times per week. ABT-263 was administered by oral gavage at a dose of 100 mg/kg BW for five consecutive days per week. ABT-263 was dissolved in a vehicle consisting of ethanol (Fujifilm, 054-07225), polyethylene glycol 400 (BioUltra, 91893-250ML-F), and Phosal 50 PG (MedChem Express, HY-H1903) at a ratio of 10:30:60.

**Evaluation of arthritis**

Joint swelling was scored ranging from 0 to 4 per paw, with a maximal score of 16 per mouse. We judged the development of arthritis in the joints using the following criteria: 0, normal; 1, mild redness, slight swelling of the ankle or wrist; 2, moderate swelling of the ankle or wrist; 3, severe swelling, including some digits, ankles or feet; 4, maximally inflamed(*32*).

**Histomorphometric analysis**

H&E and Safranin O staining of joint tissue sections were performed according to standard protocols. Ankle joints isolated from mice subjected to CAIA were fixed in 4 % paraformaldehyde at 4 °C overnight, decalcified in Osteosoft (Merck, 101728) for 12 days at 22-24 °C, embedded in paraffin, and sectioned at a thickness of 4-6 µm. H&E staining was used to assess inflammatory cell infiltration and synovial pannus formation, whereas Safranin O staining (WALDECK, 1B-463) was performed to evaluate cartilage damage. Histopathological scoring was conducted in accordance with the SMASH recommendations(*33*). Briefly, arthritis severity was evaluated across four parameters: synovial inflammation, bone erosion, proteoglycan loss, and cartilage erosion. Each parameter was scored on a scale from 0 (normal) to 3 (severe), with increments of 0.25. All histomorphometric analyses were performed in a blinded manner.

**Immunohistochemistry**

Paraffin-embedded joint tissue sections (4-6 µm thickness) were used for histological and immunofluorescence analyses. For immunofluorescence staining, sections were incubated with primary antibodies against mCherry (1:250 dilution, Abcam, ab205402) or podoplanin (1:100 dilution, clone: eBio8.1.1, Invitrogen, 14-5381-82) followed by species-appropriate fluorescent secondary antibodies. Alexa Fluor 647-conjugated anti-chicken IgY (1:500 dilution, Thermo Fisher Scientific, A32933) or Alexa Fluor 488-conjugated anti-Syrian hamster IgG (1:500 dilution, Thermo Fisher Scientific, A21110) were used as secondary antibodies. Nuclei were counterstained with Hoechst 33342 (1:3,000 dilution, Thermo Fisher Scientific, 62249). Fluorescence images were acquired using a BZ-X800 fluorescence microscope (Keyence).

**Micro-computed tomography**

Three-dimensional micro–computed tomography (micro-CT) analysis was performed on ankle joints isolated from mice subjected to CAIA. Computed tomography scanning was conducted using a TMD1000 scanner (Yamato Scientific). Three-dimensional microstructural images were reconstructed, and quantitative structural parameters were calculated using TRI/3D-BON-FCS64 software (RATOC).**Supplemental figure captions**

### Supplemental figures

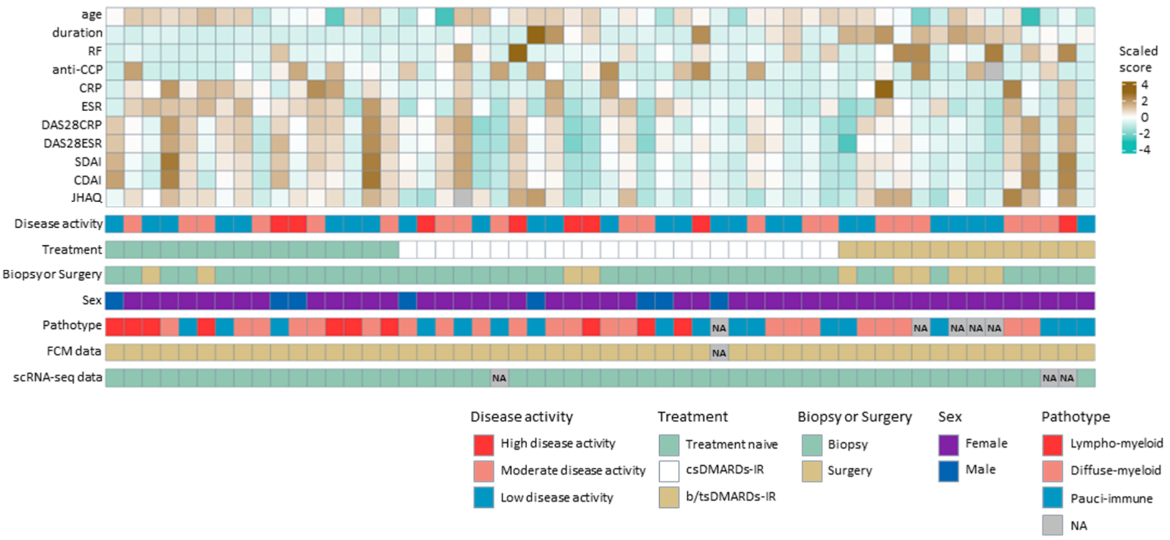


**supplemental figure S1. Clinical characteristics of patients with rheumatoid arthritis at the time of sample collection.**

Heatmap showing clinical parameters and synovial pathotypes across patients. Color scale, z-scores of continuous variables.

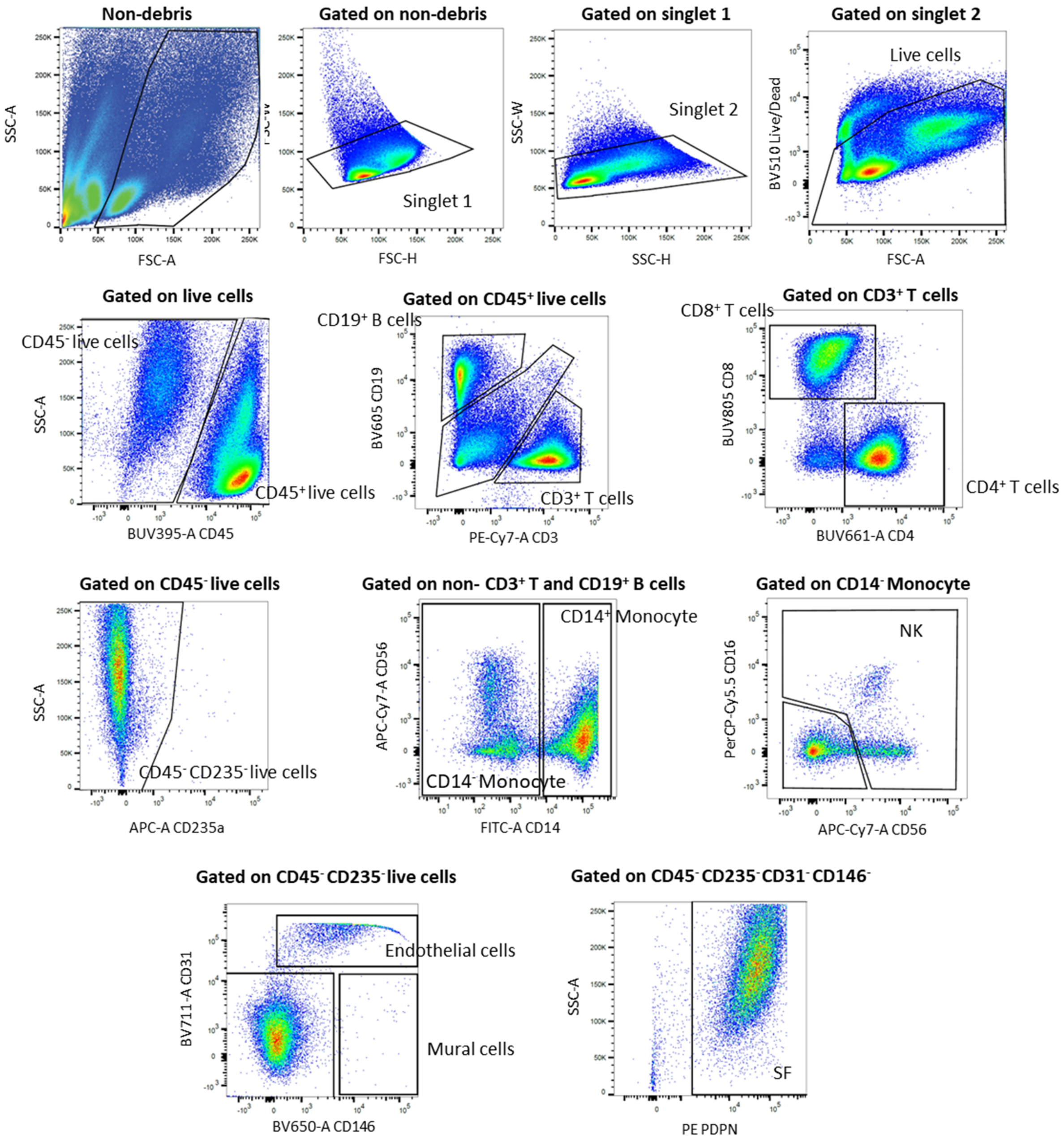


**supplemental figure S2. Gating strategy for flow cytometry analysis of synovial tissue.**

Representative flow cytometry plots showing the gating strategies used to identify $CD3^+$ T cells, $CD19^+$ B cells, monocytes, synovial fibroblasts (SF), and endothelial cells (EC). Antibodies used for flow cytometry are listed in supplemental table 10.

NK natural killer cells; SF, synovial fibroblast.

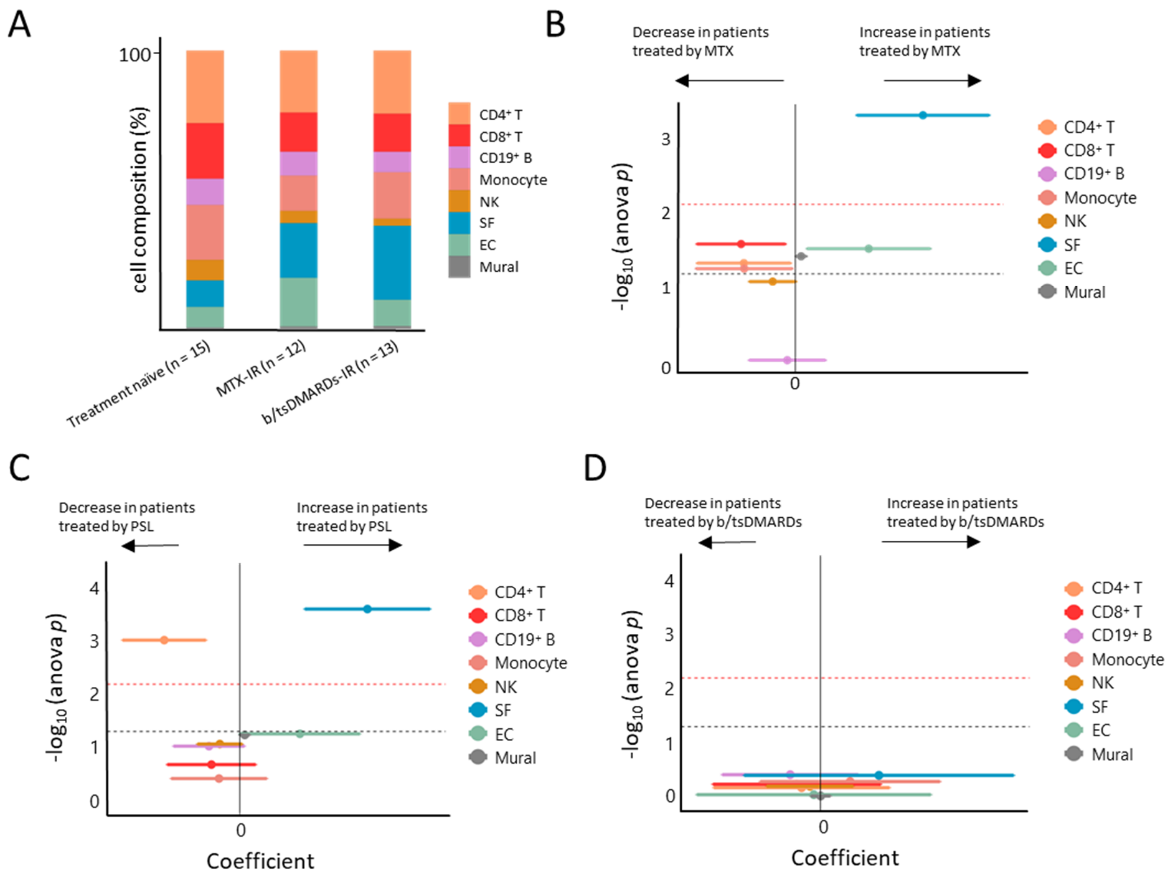


**supplemental figure S3. Dynamic changes in synovial cell populations between treatment status.**

**(A)** Synovial tissue cellular composition assessed by flow cytometry (FCM) across treatment groups: treatment-naive, methotrexate (MTX)–inadequate responders (IR), and biologic/targeted synthetic disease-modifying antirheumatic drug (b/tsDMARD)-IR patients.

**(B to D)** Association between the abundance of each synovial cell type and medication use, including MTX (**B**), prednisolone (PSL) (**C**), and b/tsDMARDs (**D**). Coefficients and association *p* values were calculated using multivariate linear regression analysis, with Clinical Disease Activity Index (CDAI) and sample collection method (biopsy or surgery) included as covariates. Central dots, estimated regression coefficients; horizontal bars, 95% confidence intervals. Red and black dashed lines indicate Bonferroni-adjusted significance thresholds and $p = 0.05$, respectively.

b/tsDMARDs, biologic/targeted synthetic Disease-Modifying Antirheumatic Drugs; CDAI, Clinical Disease Activity Index; CCP, anti-cyclic citrullinated peptide antibody; CRP, C-reactive protein; csDMARDs, conventional synthetic Disease-Modifying Antirheumatic Drugs; DAS28, Disease Activity Score in 28 joints; EC, endothelial cells; ESR, erythrocyte sedimentation rate; FCM, flow cytometry; IR, inadequate responder; JHAQ, Japanese version of the Health Assessment Questionnaire; MTX, methotrexate; NA, not applicable; NK, natural killer cells; PSL, prednisolone; RF, rheumatoid factor; SDAI, Simplified Disease Activity Index.

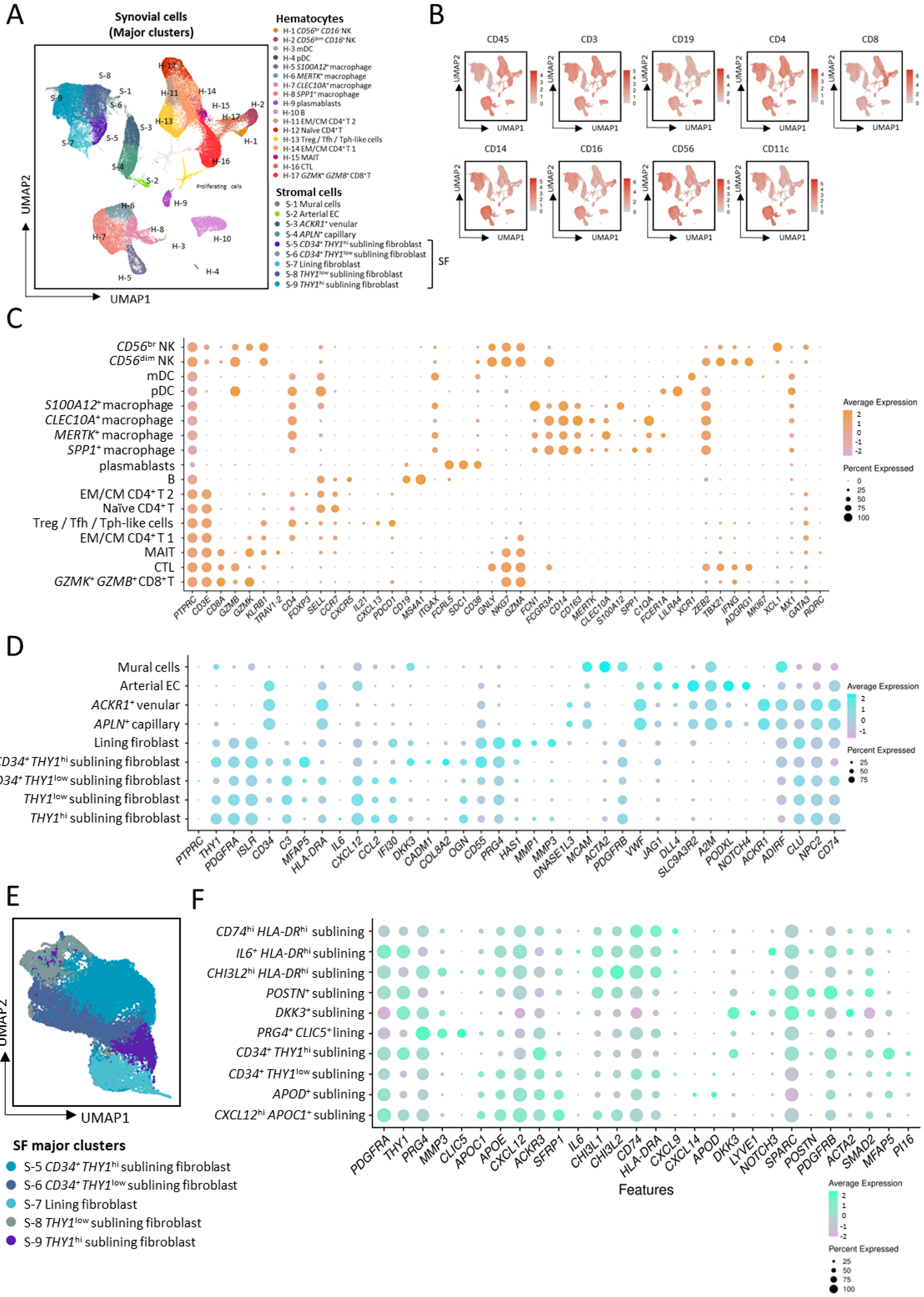

A
Synovial cells (Major clusters)
Hematocytes
H-1 CD56br CD16- NK
H-2 CD56dim CD16+ NK
H-3 mDC
H-4 pDC
H-5 S100A12+ macrophage
H-6 MERTK+ macrophage
H-7 CLEC10A+ macrophage
H-8 SPP1+ macrophage
H-9 plasmablasts
H-10 B
H-11 EM/CM CD4+ T 2
H-12 Naïve CD4+ T
H-13 Treg / Tfh / Tph-like cells
H-14 EM/CM CD4+ T 1
H-15 MAIT
H-16 CTL
H-17 GZMK+ GZMB+ CD8+ T
Stromal cells
S-1 Mural cells
S-2 Arterial EC
S-3 ACKR1+ venular
S-4 APLN+ capillary
S-5 CD34+ THY1hi sublining fibroblast
S-6 CD34+ THY1low sublining fibroblast
S-7 Lining fibroblast
S-8 THY1low sublining fibroblast
S-9 THY1hi sublining fibroblast
SF
Proliferating cells
UMAP1
UMAP2
B
CD45
CD3
CD19
CD4
CD8
CD14
CD16
CD56
CD11c
C
CD56br NK
CD56dim NK
mDC
pDC
S100A12+ macrophage
CLEC10A+ macrophage
MERTK+ macrophage
SPP1+ macrophage
plasmablasts
B
EM/CM CD4+ T 2
Naïve CD4+ T
Treg / Tfh / Tph-like cells
EM/CM CD4+ T 1
MAIT
CTL
GZMK+ GZMB+ CD8+ T
PTPRC CD3E CD8A GZMB GZMK KLRB1 TRAV1-2 CD4 FOXP3 SELL CCR7 CXCR5 IL21 CXCL13 PDCD1 CD19 MS4A1 ITGAX FCRL5 SDC1 CD38 GNLY NKG7 GZMA FCN1 FCGR3A CD14 CD163 MERTK CLEC10A S100A12 SPP1 C1QA FCER1A LILRA4 XCR1 ZEB2 TBX21 IFNG ADGRG1 MKI67 XCL1 MX1 GATA3 RORC
Average Expression
Percent Expressed
D
Mural cells
Arterial EC
ACKR1+ venular
APLN+ capillary
Lining firoblast
CD34+ THY1hi sublining fibroblast
CD34+ THY1low sublining fibroblast
THY1low sublining fibroblast
THY1hi sublining fibroblast
PTPRC THY1 PDGFRA ISLR CD34 C3 MFAP5 HLA-DRA IL6 CXCL12 CCL2 IFI30 DKK3 CADM1 COL8A2 OGN CD55 PRG4 HAS1 MMP1 MMP3 DNASE1L3 MCAM ACTA2 PDGFRB VWF JAG1 DLL4 SLC9A3R2 A2M PODXL NOTCH4 ACKR1 ADIRF CLU NPC2 CD74
Average Expression
Percent Expressed
E
UMAP1
UMAP2
SF major clusters
S-5 CD34+ THY1hi sublining fibroblast
S-6 CD34+ THY1low sublining fibroblast
S-7 Lining fibroblast
S-8 THY1low sublining fibroblast
S-9 THY1hi sublining fibroblast
F
CD74hi HLA-DRhi sublining
IL6+ HLA-DRhi sublining
CHI3L2hi HLA-DRhi sublining
POSTN+ sublining
DKK3+ sublining
PRG4+ CLIC5+ lining
CD34+ THY1hi sublining
CD34+ THY1low sublining
APOD+ sublining
CXCL12hi APOC1+ sublining
PDGFRA THY1 PRG4 MMP3 CLIC5 APOC1 APOE CXCL12 ACKR3 SFRP1 IL6 CHI3L1 CHI3L2 CD74 HLA-DRA CXCL9 CXCL14 APOD DKK3 LYVE1 NOTCH3 SPARC POSTN PDGFRB ACTA2 SMAD2 MFAP5 PI16
Features
Average Expression
Percent Expressed

**supplemental figure S4. Differential protein and gene expression across synovial cell clusters.**

**(A)** Uniform manifold approximation and projection (UMAP) integrating single-cell transcriptomic and proteomic data, showing 26 major synovial cell populations.

**(B)** Feature plot showing protein expression levels of synovial cells assessed by Cellular Indexing of Transcriptomes and Epitopes by sequencing (CITE-seq).

**(C and D)** Dot plots showing expression of representative marker proteins in major hematopoietic ($CD45^{+}$) (**C**) and stromal ($CD45^{-}$) (**D**) synovial cell clusters.

**(E)** Major synovial fibroblast (SF) clusters projected onto uniform manifold approximation and projection (UMAP) and annotated by fine clusters.

**(F)** Dot plot showing gene expression patterns across SF fine clusters.

CITE-seq, Cellular Indexing of Transcriptomes and Epitopes by sequencing; CTL, cytotoxic T lymphocyte; EC, endothelial cell; MAIT, mucosal-associated invariant T cells; mDC, myeloid dendritic cell; NK, natural killer cell; pDC, plasmacytoid dendritic cell; SF, synovial fibroblast; Tfh, follicular helper T cell; Tph, peripheral helper T cell; Treg, regulatory T cell; UMAP, uniform manifold approximation and projection.

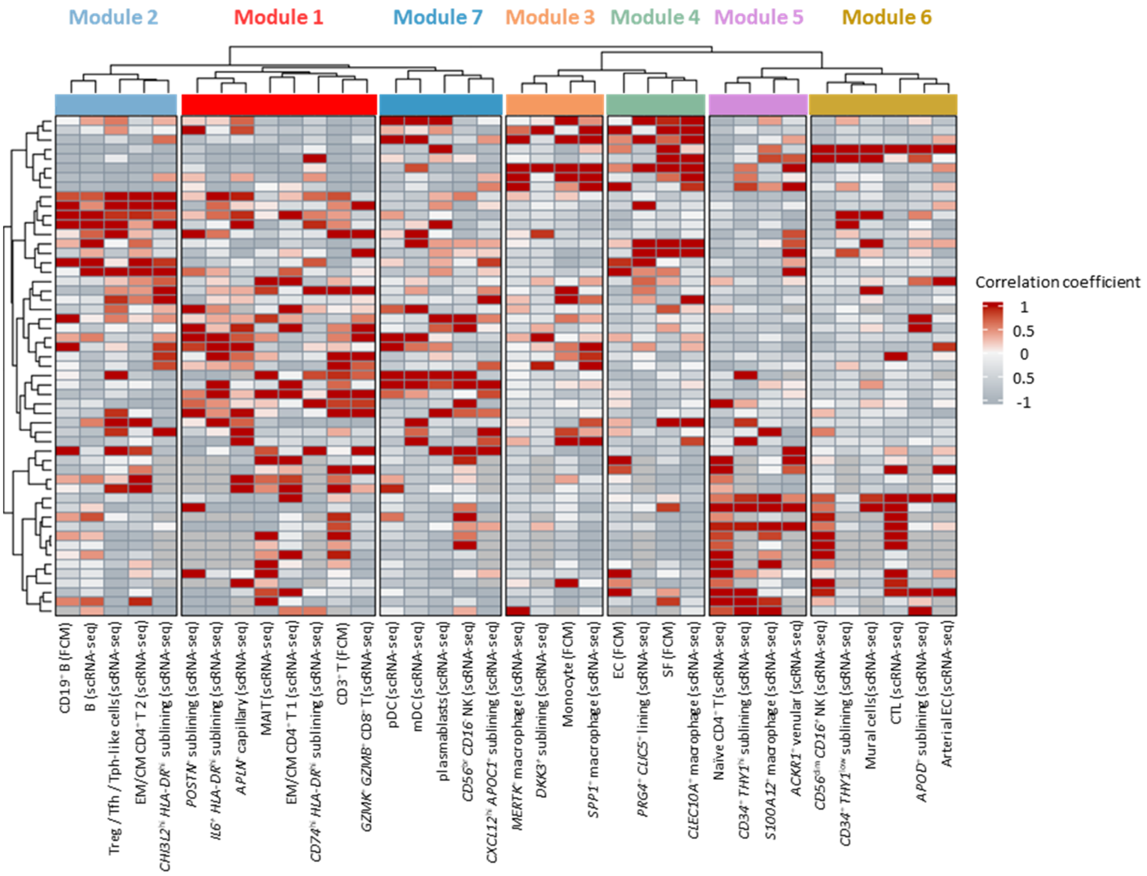


**supplemental figure S5. Hierarchical clustering identifies seven synovial cell modules.**

Hierarchical clustering based on the relative abundance of synovial cell types quantified by flow cytometry ($CD3^+$ T cells, $CD19^+$ B cells, monocytes, synovial fibroblasts (SF), and endothelial cells, expressed as a percentage of total live cells) and single-cell RNA sequencing (major hematopoietic cell clusters as a percentage of total hematopoietic cells, major non-SF stromal cell clusters as a percentage of total non-SF stromal cells, and SF fine clusters as a percentage of total SF). Columns, cell types and clusters; rows, individual subjects.

CTL, cytotoxic T lymphocyte; EC, endothelial cell; FCM, flow cytometry; MAIT, mucosal-associated invariant T cells; mDC, myeloid dendritic cell; NK, natural killer cell; pDC, plasmacytoid dendritic cell; scRNA-seq, single cell RNA sequencing; SF, synovial fibroblast; Tfh, follicular helper T cell; Tph, peripheral helper T cell; Treg, regulatory T cell.

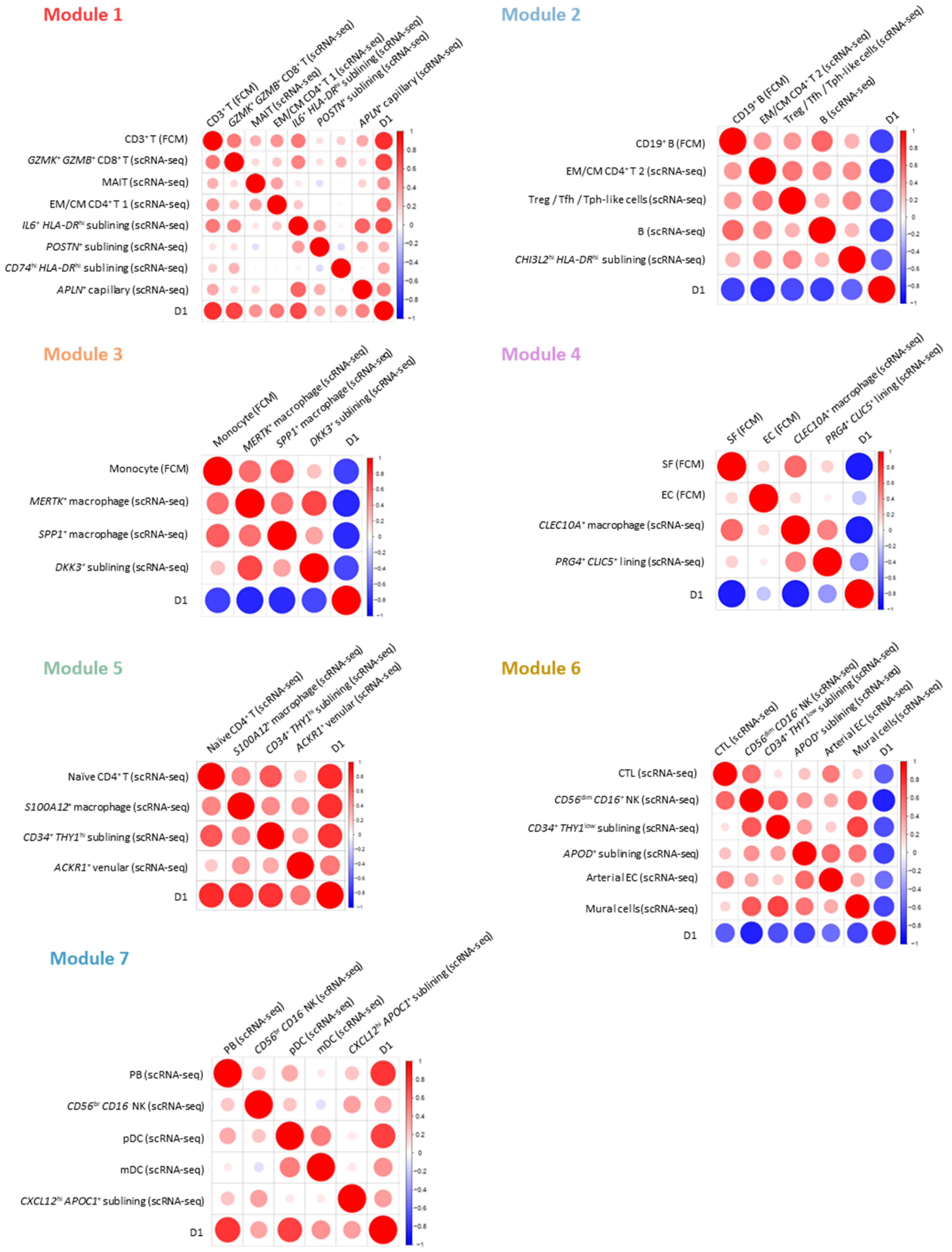


**supplemental figure S6. Definition of the MOSAIC score by multidimensional scaling.**

Correlation plot showing Pearson's correlation coefficients between the proportion of each cell cluster and the first dimension (D1) obtained by multidimensional scaling for each cell module. The MOdular Stromal And Immune Cells abundance (MOSAIC) score corresponds to the D1 score. When D1 is negatively correlated with the cell proportions, the MOSAIC score is defined as the sign-inverted D1 value.

CTL, cytotoxic T lymphocyte; EC, endothelial cell; FCM, flow cytometry; MAIT, mucosal-associated invariant T cells; mDC, myeloid dendritic cell; MOSAIC, MOdular Stromal And Immune Cells abundance; NK, natural killer cell; pDC, plasmacytoid dendritic cell; scRNA-seq, single cell RNA sequencing; SF, synovial fibroblast; Tfh, follicular helper T cell; Tph, peripheral helper T cell; Treg, regulatory T cell.

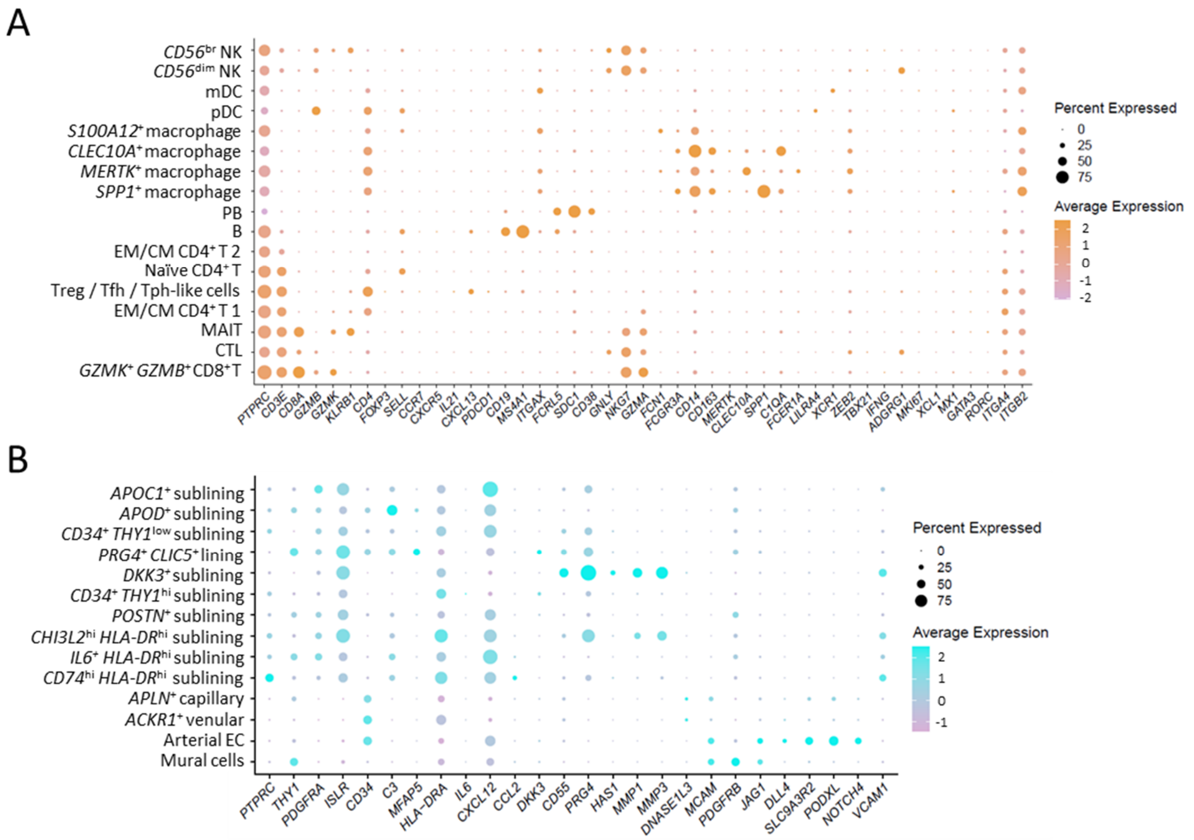


**supplemental figure S7. Transcriptomic characteristics of synovial fibroblast fine clusters defined by Xenium and snATAC-seq.**

**(A and B)** Gene expression profiles of **(A)** hematocytes (CD45$^{+}$ cells) and **(B)** stromal cells (CD45$^{-}$ cells) identified by spatial transcriptomics (Xenium, 10x Genomics) through reference mapping onto a single-cell RNA-sequencing (scRNA-seq) dataset.

scRNA-seq, single-cell RNA sequencing.

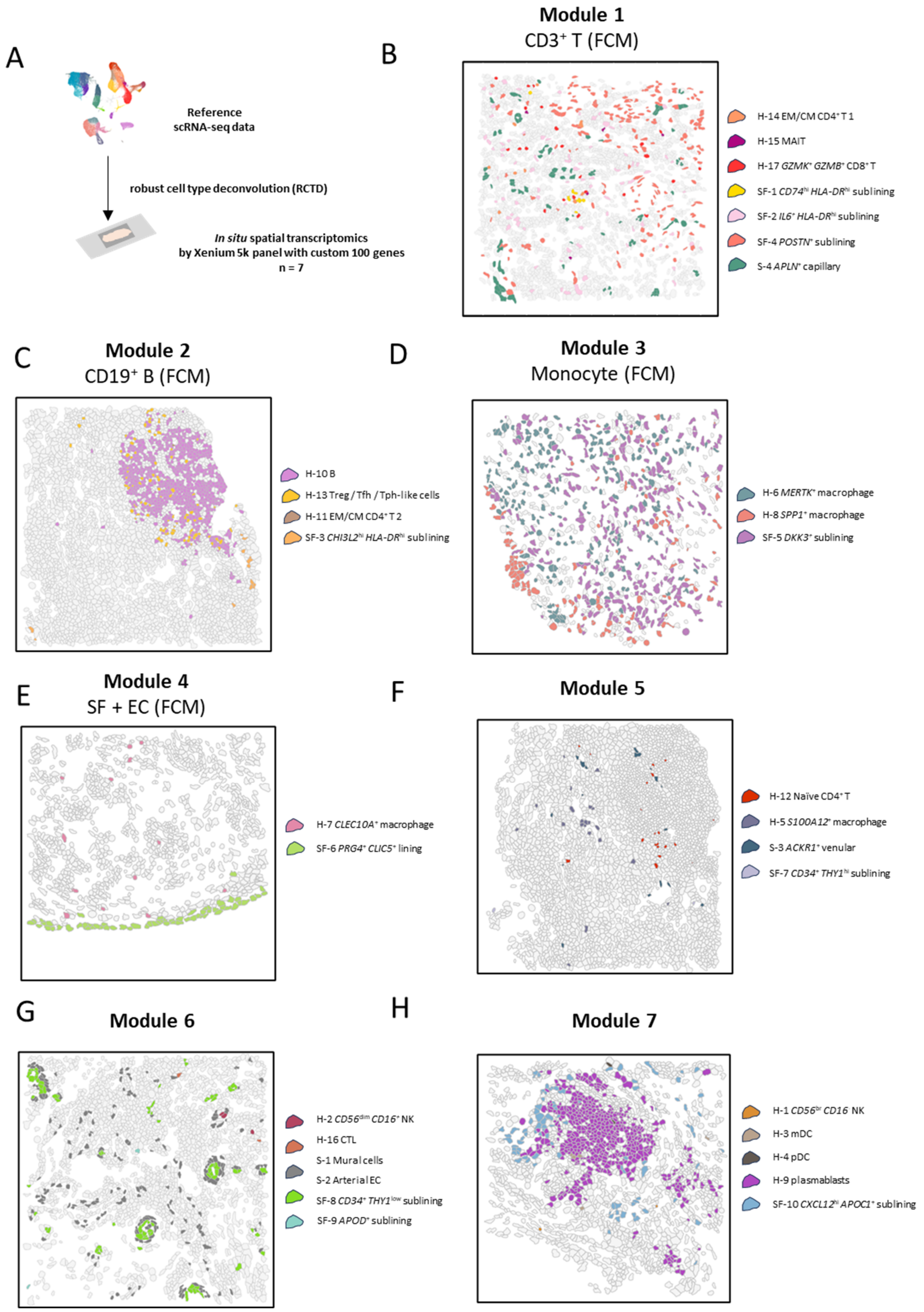

A
Reference scRNA-seq data
robust cell type deconvolution (RCTD)
In situ spatial transcriptomics by Xenium 5k panel with custom 100 genes
n = 7
B
Module 1
CD3+ T (FCM)
H-14 EM/CM CD4+ T 1
H-15 MAIT
H-17 GZMK+ GZMB+ CD8+ T
SF-1 CD74hi HLA-DRhi sublining
SF-2 IL6+ HLA-DRhi sublining
SF-4 POSTN+ sublining
S-4 APLN+ capillary
C
Module 2
CD19+ B (FCM)
H-10 B
H-13 Treg / Tfh / Tph-like cells
H-11 EM/CM CD4+ T 2
SF-3 CHI3L2hi HLA-DRhi sublining
D
Module 3
Monocyte (FCM)
H-6 MERTK+ macrophage
H-8 SPP1+ macrophage
SF-5 DKK3+ sublining
E
Module 4
SF + EC (FCM)
H-7 CLEC10A+ macrophage
SF-6 PRG4+ CLIC5+ lining
F
Module 5
H-12 Naïve CD4+ T
H-5 S100A12+ macrophage
S-3 ACKR1+ venular
SF-7 CD34+ THY1hi sublining
G
Module 6
H-2 CD56dim CD16+ NK
H-16 CTL
S-1 Mural cells
S-2 Arterial EC
SF-8 CD34+ THY1low sublining
SF-9 APOD+ sublining
H
Module 7
H-1 CD56br CD16- NK
H-3 mDC
H-4 pDC
H-9 plasmablasts
SF-10 CXCL12hi APOC1+ sublining

**supplemental figure S8. Spatial organization of seven cell modules underlying the MOdular Stromal And Immune Cells (MOSAIC) abundance.**

**(A)** Schematic overview illustrating the reference-mapping workflow used to annotate cells in the spatial data by transferring labels from our reference scRNA-seq dataset using robust cell type deconvolution (RCTD).

**(B to H)** Representative images showing the spatial distribution of cells assigned to each cell module within synovial tissue sections.

CTL, cytotoxic T lymphocyte; EC, endothelial cell; FCM, flow cytometry; MAIT, mucosal-associated invariant T cells; mDC, myeloid dendritic cell; MOSAIC, MOdular Stromal And Immune Cells abundance; NK, natural killer cell; pDC, plasmacytoid dendritic cell; RCTD, Robust Cell Type Decomposition; scRNA-seq, single cell RNA sequencing; SF, synovial fibroblast; Tfh, follicular helper T cell; Tph, peripheral helper T cell; Treg, regulatory T cell.

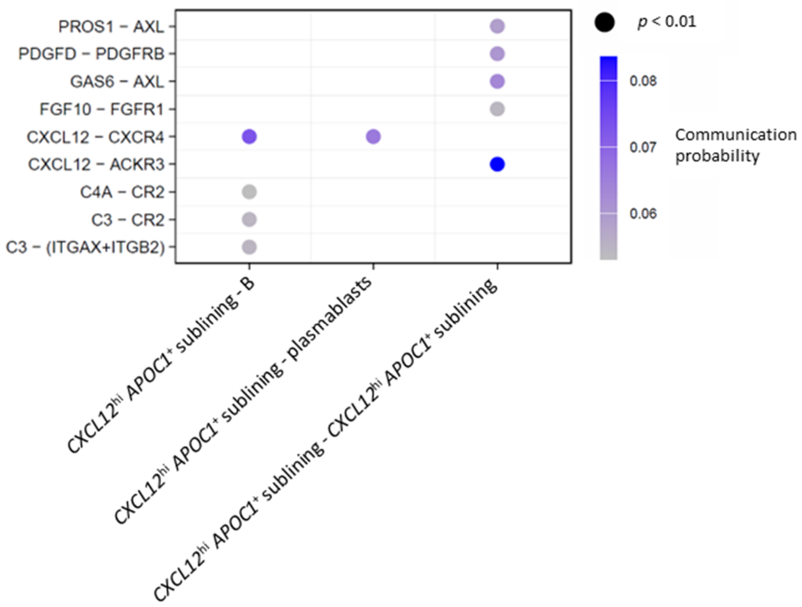


**supplemental figure S9. Intercellular communication is dominated by CXCL12-associated signaling pathways involving *CXCL12*$^{hi}$ *APOC1*$^{+}$ sublining fibroblasts and plasmablasts.**

Bubble plot visualizing ligand-receptor interactions mediated by CXCL12-associated pathways among *CXCL12*$^{hi}$ *APOC1*$^{+}$ sublining synovial fibroblasts (SF), B cells and plasmablasts.

SF, synovial fibroblast.

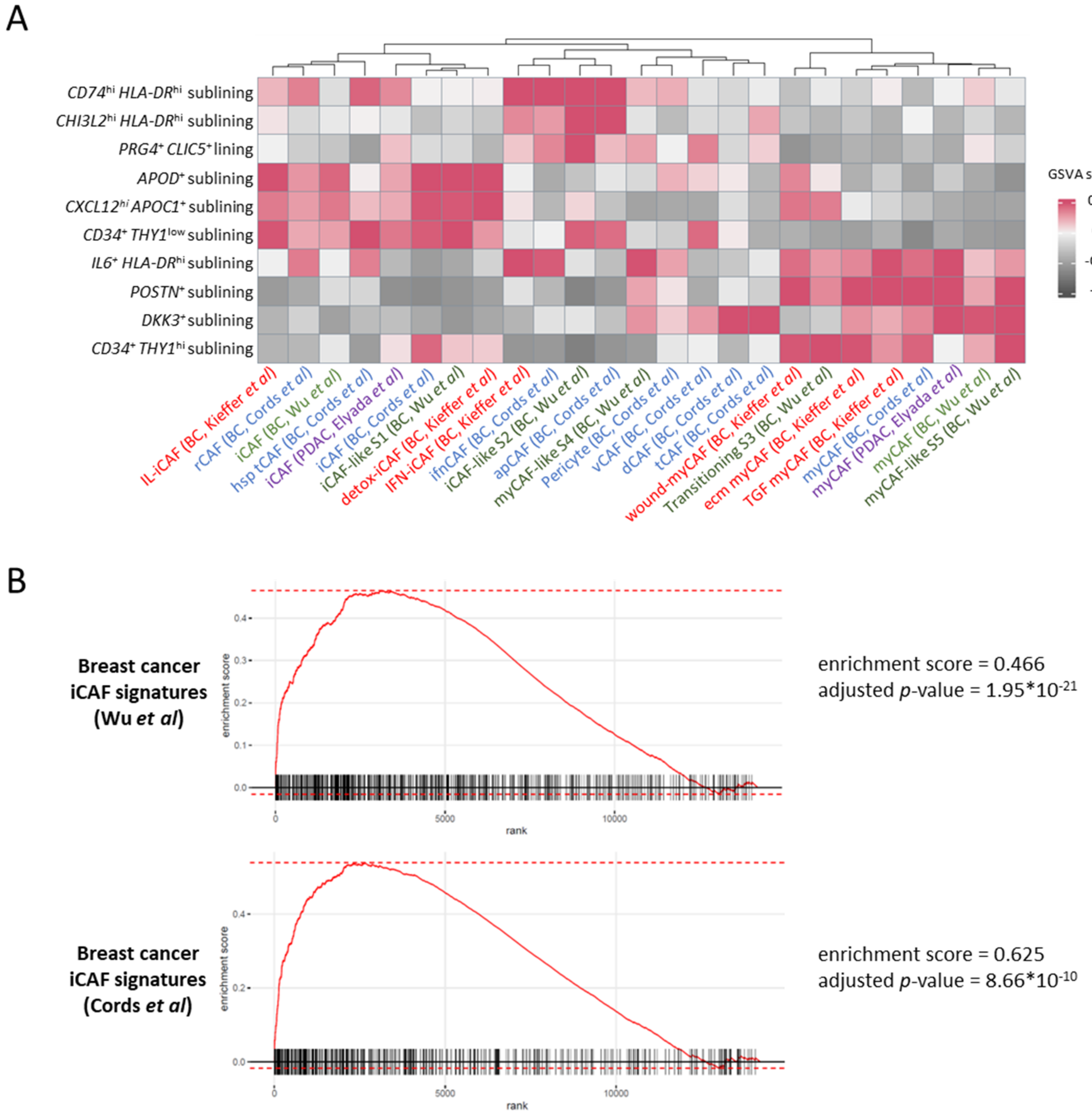


**supplemental figure S10. *CXCL12*$^{hi}$ *APOC1*$^{+}$ sublining fibroblasts share a common transcriptional program with inflammatory cancer-associated fibroblasts (CAF).**

**(A)** Heatmap showing Gene Set Variation Analysis (GSVA) scores for selected gene signatures derived from published CAF populations from breast and pancreatic cancers across SF fine clusters.

**(B)** Enrichment of inflammatory CAF (iCAF) signatures derived from published breast cancer CAF datasets in negative control relative to *APOC1*-depleted SF, calculated by Gene Set Enrichment Analysis (GSEA).

CAF, cancer-associated fibroblast; GSEA, gene set enrichment analysis; GSVA, gene set variation analysis; SF, synovial fibroblast.

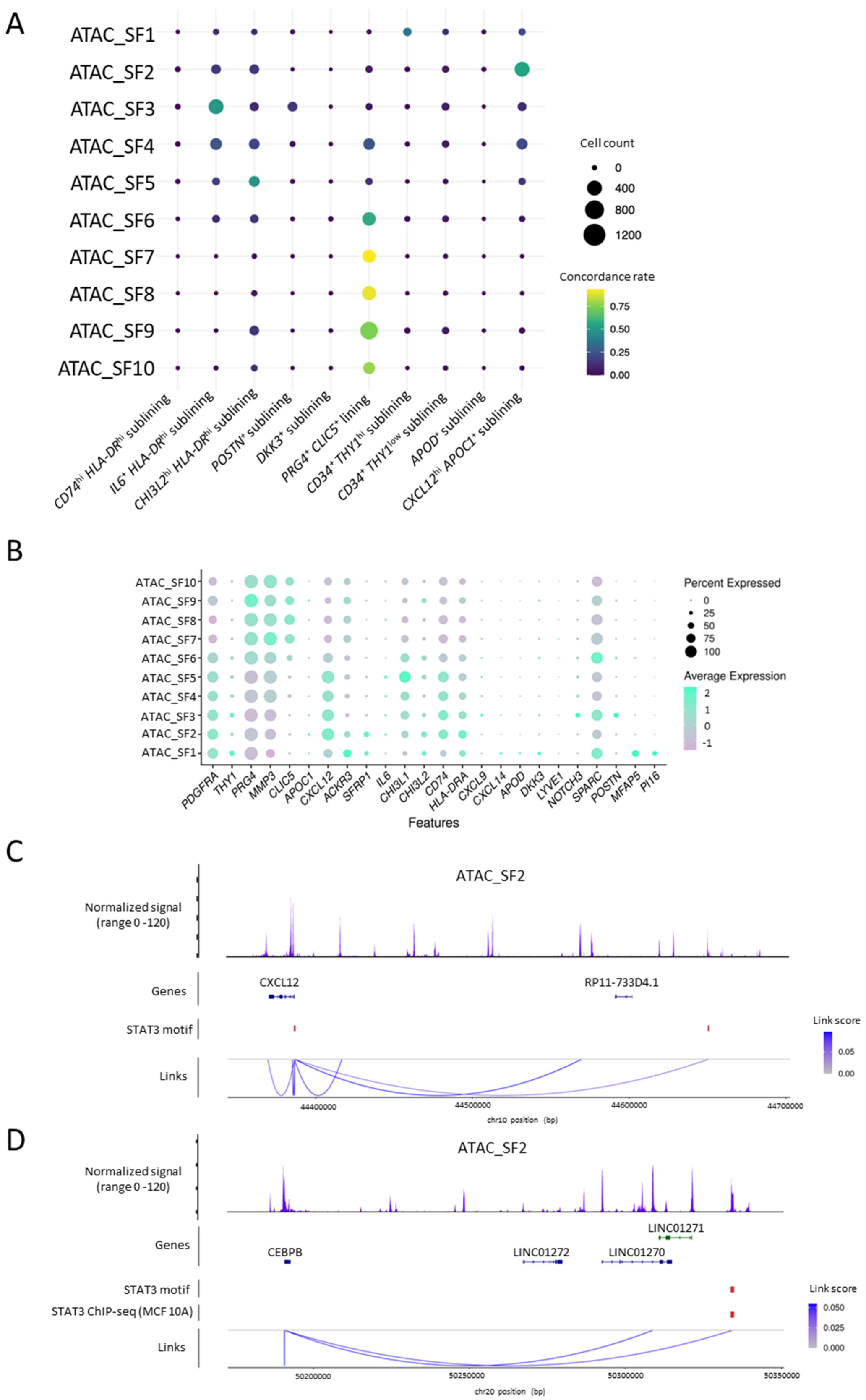

A
ATAC_SF1
ATAC_SF2
ATAC_SF3
ATAC_SF4
ATAC_SF5
ATAC_SF6
ATAC_SF7
ATAC_SF8
ATAC_SF9
ATAC_SF10
CD74hi HLA-DRhi sublining
IL6+ HLA-DRhi sublining
CHI3L2hi HLA-DRhi sublining
POSTN+ sublining
DKK3+ sublining
PRG4+ CLIC5+ lining
CD34+ THY1hi sublining
CD34+ THY1low sublining
APOD+ sublining
CXCL12hi APOC1+ sublining
Cell count
0
400
800
1200
Concordance rate
0.75
0.50
0.25
0.00
B
Percent Expressed
0
25
50
75
100
Average Expression
2
1
0
-1
PDGFRA
THY1
PRG4
MMP3
CLIC5
APOC1
CXCL12
ACKR3
SFRP1
IL6
CHI3L1
CHI3L2
CD74
HLA-DRA
CXCL9
CXCL14
APOD
DKK3
LYVE1
NOTCH3
SPARC
POSTN
MFAP5
PI16
Features
C
ATAC_SF2
Normalized signal
(range 0 -120)
Genes
CXCL12
RP11-733D4.1
STAT3 motif
Links
Link score
0.05
0.00
44400000
44500000
44600000
44700000
chr10 position (bp)
D
ATAC_SF2
Normalized signal
(range 0 -120)
Genes
LINC01271
CEBPB
LINC01272
LINC01270
STAT3 motif
STAT3 ChIP-seq (MCF 10A)
Links
Link score
0.050
0.025
0.000
50200000
50250000
50300000
50350000
chr20 position (bp)

**supplemental figure S11. Identification of the transcriptional machinery characterizing *CXCL12*$^{hi}$ *APOC1*$^{+}$ fibroblasts.**

**(A)** Concordance between SF fine clusters defined by single-nucleus RNA sequencing combined with assay for transposase-accessible chromatin using sequencing (snRNA-seq + snATAC-seq) and those defined by single-cell RNA sequencing (scRNA-seq). The snRNA-seq + snATAC-seq dataset was mapped onto the scRNA-seq reference using RNA expression profiles with Symphony. Bubble size, the number of cells assigned to each pair of clusters; color scale, the proportion of cells assigned to each scRNA-seq cluster among SF fine clusters defined by chromatin accessibility.

**(B)** Nuclear RNA expression of marker genes in synovial fibroblast (SF) fine clusters defined by single-nucleus assay for transposase-accessible chromatin sequencing (snATAC-seq).

**(C and D)** Organization of transcriptional regulatory regions surrounding *CXCL12* **(C)** and *C/EBPB* **(D)** in ATAC_SF2. Link color intensity represents the link score, defined as the correlation coefficient between gene expression and accessibility of individual peaks located within ±500 kb of the transcription starting site (TSS). Highlighted peaks include those containing *STAT3* motifs (link score > 0.05, $p < 0.05$) and overlapping with public STAT3 ChIP-seq peaks from MCF10A cells (SRX150479, SRX150536, SRX150630, SRX150670, SRX4192887, SRX10119380, SRX10119382). *P* values correspond to the z-score-based significance of correlation coefficients relative to background peaks.

ChIP-seq, Chromatin immunoprecipitation sequencing; scRNA-seq, single-cell RNA sequencing; SF, synovial fibroblast; snATAC-seq, single-nucleus assay for transposase-accessible chromatin sequencing; snRNA-seq, single-nucleus RNA sequencing; TSS, transcription starting site; UMAP, uniform manifold approximation and projection.

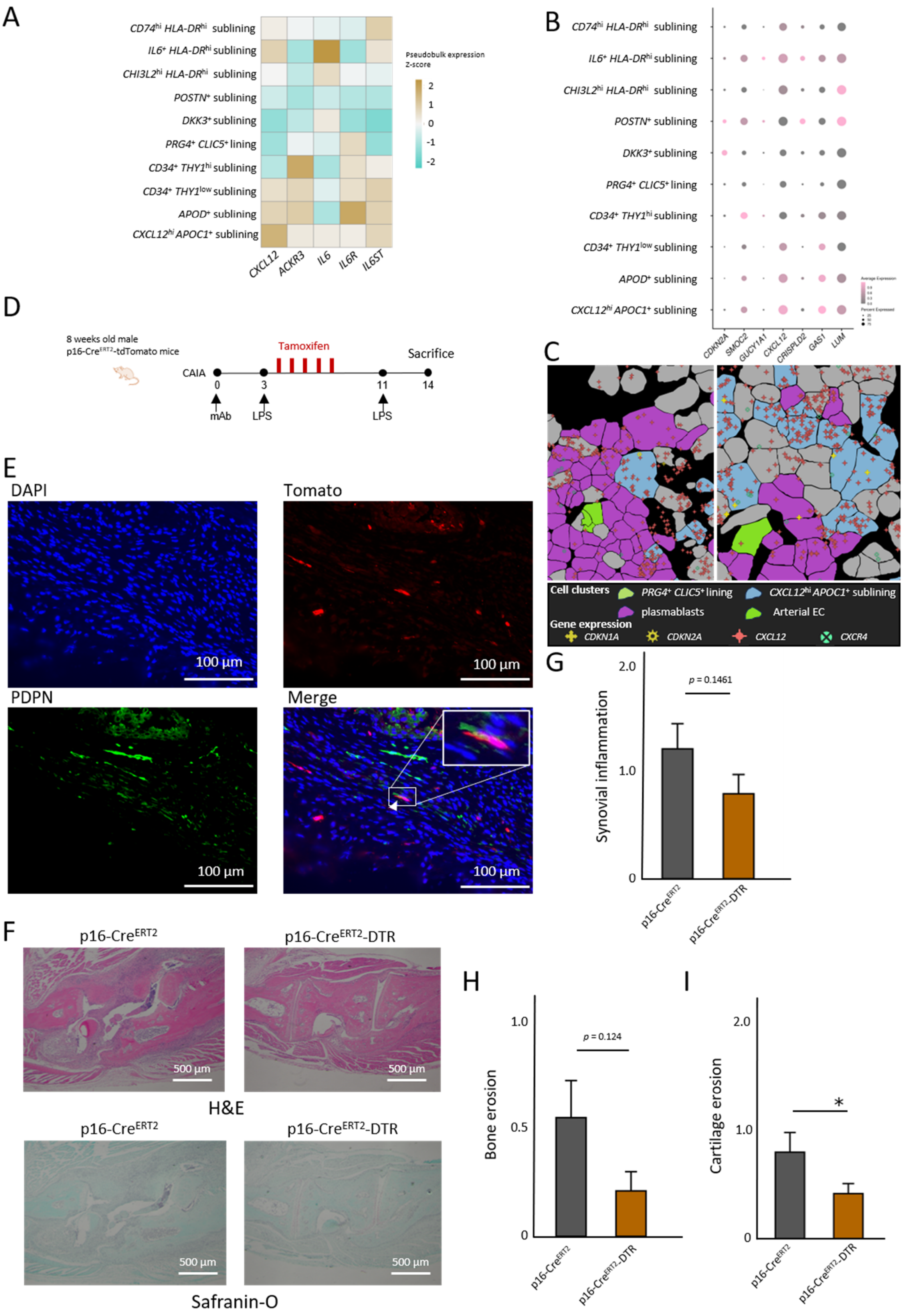
A
CD74hi HLA-DRhi sublining
IL6+ HLA-DRhi sublining
CHI3L2hi HLA-DRhi sublining
POSTN+ sublining
DKK3+ sublining
PRG4+ CLIC5+ lining
CD34+ THY1hi sublining
CD34+ THY1low sublining
APOD+ sublining
CXCL12hi APOC1+ sublining
CXCL12
ACKR3
IL6
IL6R
IL6ST
Pseudobulk expression
Z-score
2
1
0
-1
-2
B
CDKN2A
SMOC2
GUCY1A1
CXCL12
CRISPLD2
GAS1
LUM
C
Cell clusters
PRG4+ CLIC5+ lining
CXCL12hi APOC1+ sublining
plasmablasts
Arterial EC
Gene expression
CDKN1A
CDKN2A
CXCL12
CXCR4
D
8 weeks old male
p16-CreERT2-tdTomato mice
Tamoxifen
Sacrifice
CAIA
0
3
11
14
mAb
LPS
LPS
E
DAPI
Tomato
PDPN
Merge
100 µm
G
Synovial inflammation
p = 0.1461
2.0
1.0
0
p16-CreERT2
p16-CreERT2-DTR
F
p16-CreERT2
p16-CreERT2-DTR
500 µm
H&E
Safranin-O
H
Bone erosion
p = 0.124
1.0
0.5
0
I
Cartilage erosion
*
2.0
1.0
0

**supplemental figure S12. Presence of p16$^{h}$-sn fibroblasts in synovial tissue and the effect of prophylactic elimination of p16$^{h}$-sn cells on experimental arthritis.**

**(A)** Heatmap depicting the cluster-level average expression of SASP factors and their receptors across SF fine clusters, scaled per gene across clusters.

**(B)** Dot plot showing expression of marker genes defining the p16$^{h}$-sn cancer-associated fibroblasts (CAF) from Meguro *et al* across synovial fibroblasts (SF) fine clusters.

**(C)** Representative images showing *CXCL12*$^{hi}$ *APOC1*$^{+}$ fibroblasts expressing *CDKN1A* or *CDKN2A* surrounded by plasmablasts in synovial tissue from a patient with active rheumatoid arthritis, together with *CXCL12* and *CXCR4* expression.

**(D)** Experimental design of the collagen antibody-induced arthritis (CAIA) mouse model using 8-week-old male p16-Cre$^{ERT2}$-tdTomato mice with tamoxifen administration. Following the lipopolysaccharide (LPS) boost, mice received intraperitoneal injections of tamoxifen (80 mg/kg) for five consecutive days.

**(E)** Representative immunofluorescence images of inflamed synovial tissue from p16-Cre$^{ERT2}$-tdTomato mice.

**(F)** Representative hematoxylin and eosin (H&E) and Safranin O-stained images of right ankle joints from p16-Cre$^{ERT2}$-DTR mice and p16-Cre$^{ERT2}$ control mice under CAIA conditions.

**(G to I)** Histopathological scores for synovial inflammation (**e**), bone erosion (**f**), and cartilage destruction (**g**) in p16-Cre$^{ERT2}$-DTR mice (n = 12) and p16-Cre$^{ERT2}$ control mice (n = 12) under CAIA conditions. Bars, mean; error bars, SEM. *P* values, two-tailed Mann-Whitney *U* test. (* $p < 0.05$). One p16-Cre$^{ERT2}$ control mouse was excluded from the histological analysis because tissue sections could not be successfully prepared.

CAIA, collagen antibody-induced arthritis; CAF, cancer-associated fibroblast; DAPI, 4',6-diamidino-2-phenylindole; DTR, diphtheria toxin receptor; H&E, haematoxylin and eosin; LPS, lipopolysaccharide; mAb, monoclonal antibody; PDPN, podoplanin; SASP, senescence-associated secretory phenotype; SEM, standard error of the mean; SF, synovial fibroblast; sn, senescent.

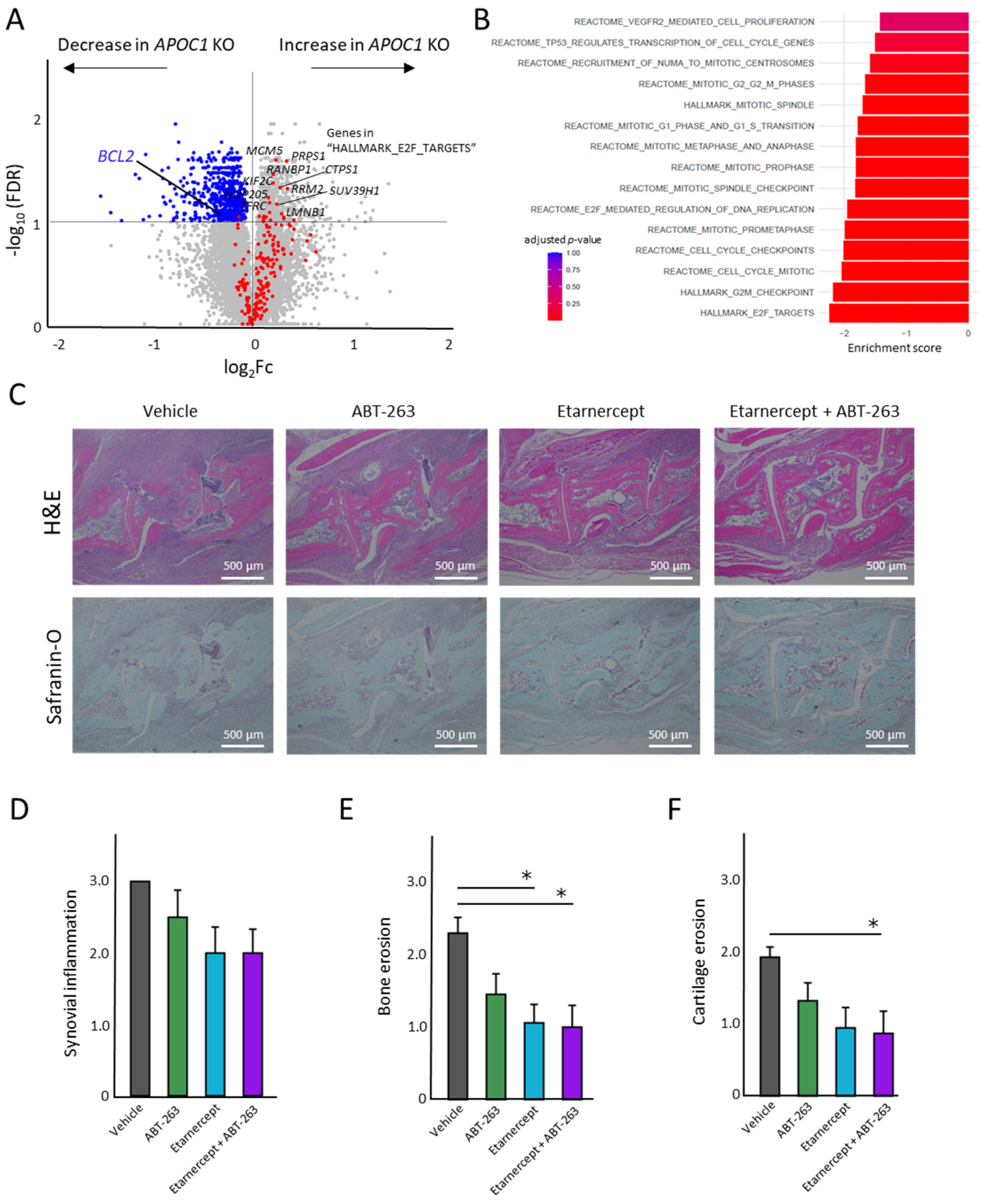


**supplemental figure S13. Combined TNF inhibition and senolytic treatment effectively reduces bone and cartilage erosion.**

**(A)** Differential gene expression analysis comparing synovial fibroblasts (SF) depleted of *APOC1* or negative control. Blue points, genes with significantly decreased expression (false discovery rate (FDR) < 0.01); blue label, *BCL2*; red dots, genes in the "HALLMARK_E2F_TARGETS" which encodes cell-cycle–related targets of E2F transcription factors from the Molecular Signatures Database (MSigDB).

**(B)** Enrichment of cell-cycle–related pathways in negative control relative to *APOC1*-depleted SF, calculated by Gene Set Enrichment Analysis (GSEA).

**(C)** Representative H&E- and Safranin O-stained images of right ankle joints from mice treated with etanercept plus ABT-263, etanercept alone, ABT-263 alone, or vehicle under CAIA conditions.

**(D to F)** Histopathological scores for synovial inflammation (**D**), bone erosion (**E**), and cartilage destruction (**F**) in mice treated with etanercept plus ABT-263 (n = 9), etanercept alone (n = 9), ABT-263 alone (n = 8), or vehicle (n = 9) under CAIA conditions. Bars, mean; error bars, SEM. *P* values, the Kruskal–Wallis rank-sum test, followed by Dunn's multiple comparison test with Holm's correction (* $p < 0.05$).

CAIA, collagen antibody-induced arthritis; DAPI, 4',6-diamidino-2-phenylindole; DTR, diphtheria toxin receptor; Fc, fold change; FDR, false discovery rate; GSEA, Gene Set Enrichment Analysis; H&E, hematoxylin and eosin; LPS, lipopolysaccharide; mAb, monoclonal antibody; PDPN, podoplanin; SEM, standard error of the mean; SF, synovial fibroblast; TNF, tumor necrosis factor.

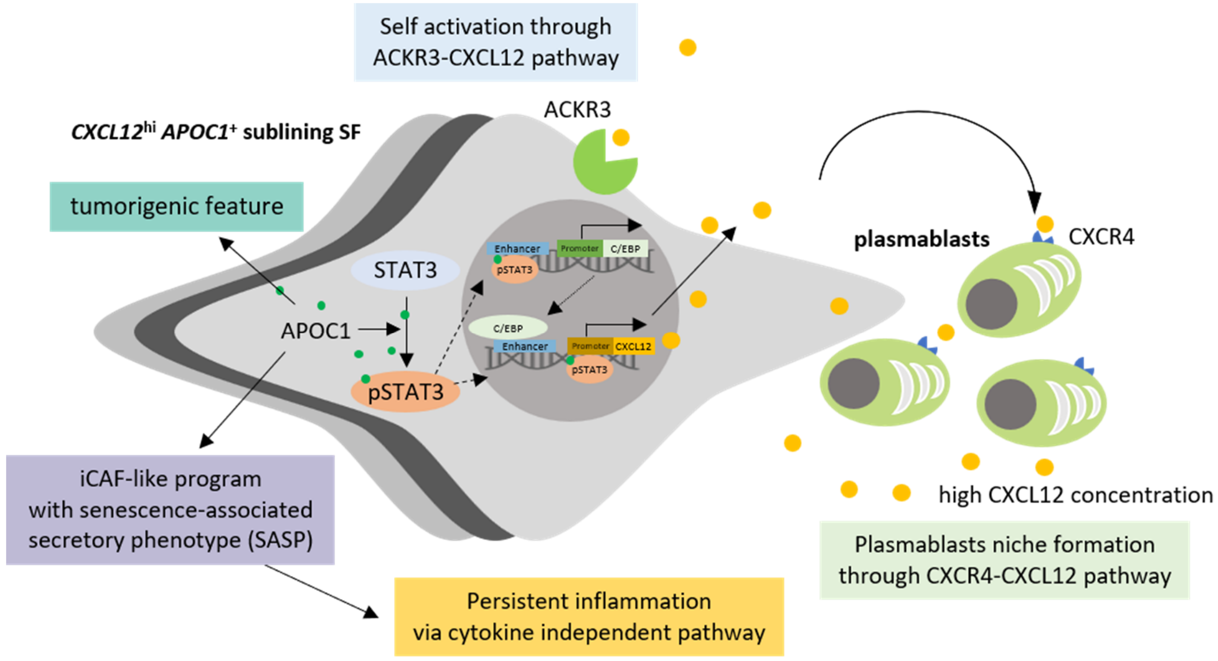

**supplemental figure S14. Proposed model of APOC1-driven pathogenic synovial fibroblast states associated with treatment resistance.**

*CXCL12*$^{hi}$ *APOC1*$^{+}$ sublining synovial fibroblasts (SF) exhibit tumor-like properties driven by APOC1 while retaining inflammatory cancer-associated fibroblast (iCAF)-like features. These cells are characterized by high CXCL12 expression, a representative senescence-associated secretory phenotype (SASP) factor, and activation of C/EBP family transcription factors. APOC1 is proposed to promote STAT3 phosphorylation, leading to transcriptional activation of C/EBP family members. In turn, C/EBP factors engage enhancer elements of *CXCL12*, while STAT3 binds its promoter, resulting in sustained CXCL12 production. By establishing a CXCL12-enriched niche within the synovium, *CXCL12*$^{hi}$ *APOC1*$^{+}$ sublining SF support plasma cell differentiation and infiltration, thereby contributing to resistance to cytokine-blocking therapies. This mechanism is analogous to the role of senescent iCAFs in promoting cancer progression through SASP production.

iCAF, inflammatory cancer-associated fibroblast; SASP, senescence-associated secretory phenotype; SF, synovial fibroblast.
